# Arrhythmia Mechanisms and Spontaneous Calcium Release: I - Multi-scale Modelling Approaches

**DOI:** 10.1101/434464

**Authors:** Michael A. Colman

## Abstract

**Motivation:** Spontaneous sub-cellular calcium release events (SCRE), controlled by microscopic stochastic fluctuations of the proteins responsible for intracellular calcium release, are conjectured to promote the initiation and perpetuation of rapid arrhythmia associated with conditions such as heart failure and atrial fibrillation: SCRE may underlie the emergence of spontaneous excitation in single cells, resulting in arrhythmic triggers in tissue. However, translation of single-cell data to the tissue scale is non-trivial due to complex substrate considerations. Computational modelling provides a viable approach to dissect these multi-scale mechanisms, yet there remains a significant challenge in accurately and efficiently modelling this probabilistic behaviour in large-scale tissue models. The aim of this study was to develop an approach to overcome this challenge.

**Methods:** The dynamics of SCRE under multiple conditions (pacing rate, beta-stimulation, disease remodelling) in a computational model of stochastic, spatio-temporal calcium handling were analysed in order to develop Spontaneous Release Functions, which capture the variability and properties of SCRE matched to the full cell model. These functions were then integrated with tissue models, comprising idealised 2D sheets as well as full reconstructions of ventricular and atrial anatomy.

**Results:** The Spontaneous Release Functions accurately reproduced the dynamics of SCRE and its dependence on environment variables under multiple different conditions observed in the full single-cell model. Differences between cellular models and conditions where enhanced at the tissue scale, where the emergence of a focal excitation is largely an all-or-nothing response. Generalisation of the approaches was demonstrated through integration with an independent cell model, and parameterisation to an experimental dataset.

**Conclusions:** A novel approach has been developed to dynamically model SCRE at the tissue scale, in-line with behaviour observed in detailed single-cell models. Such an approach allows evaluation of the potential importance of SCRE in arrhythmia in both general mechanistic and disease-specific investigation.

## Introduction

Cardiovascular disease is one of the major healthcare problems faced by the developed world, with increasing prevalence associated with aging populations [1–3]. Improved understanding of the mechanisms underlying cardiac arrhythmias, a major component of cardiovascular diseases’ impact on morbidity and mortality, is therefore vital to the effort to improve both lifespan and quality of life therein. Rapid arrhythmias, such as tachycardia and fibrillation, are associated with highly non-linear electrical excitation patterns and are challenging to manage [4,5], in part because the complex multi-scale mechanisms have yet to be full elucidated.

Malfunction of the intracellular calcium (Ca^2+^) handling system has been implicated in the development of rapid arrhythmias, linking sub-cellular spontaneous Ca^2+^ release events (SCRE, described below) to pro-arrhythmic triggers in single cell [6–10]. However, translation of these cellular data to assess the mechanisms and importance of SCRE in tissue-scale arrhythmia remains a significant challenge [11]. Computational modelling provides a viable approach for detailed multi-scale evaluation of cardiac arrhythmia mechanisms. Never-the-less, simulating SCRE in tissue-scale models is non-trivial due to the complex cellular structures and non-linear, spatio-temporal dynamics underlying the associated phenomena. These are described in detail below to facilitate understanding of the challenges involved.

The intracellular Ca^2+^ handling system is a critical component of cardiac excitation-contraction coupling through the process of Ca^2+^-induced-Ca^2+^-release (CICR) [12]: Ca^2+^ enters the cell through the L-type Ca^2+^ channels (LTCC) during the electrical action potential (AP) and triggers a large release of Ca^2+^ from the sarcoplasmic reticulum (SR), the intracellular Ca^2+^ store. This Ca^2+^ release is controlled by opening of the ryanodine receptors (RyR) which lie on the membrane of the SR in a co-localised junction with the LTCCs on the surface sarcolemma membrane and transverse-tubules (t-tubules), called a dyad (Figure 1Ai). During relaxation, SR-Ca^2+^ is restored through the SR-Ca^2+^ pump (SERCA) and extruded from the cell through the sodium-Ca^2+^ exchanger (NCX) and the membrane Ca^2+^ pump.

The distributed spatial structure of the sarcolemmal and SR membranes (Figure 1Ai) and their associated channel proteins facilitates uniform cellular contraction, but also has potentially proarrhythmic implications: Each dyad contains only a few LTCC and RyR channels (typically 5-15 and 5-200, respectively [13]) located within a very small volume (O(10^−3^) μm^3^) which functionally acts as a restricted Ca^2+^ sub-space to facilitate CICR. Random openings of single or few RyRs can thus raise the local Ca^2+^ concentration sufficiently to trigger further openings within the dyad, potentially leading to a whole-dyad event: the Ca^2+^ spark (Figure 1Ai). Spatial-diffuse coupling amplifies this intrinsic feedback mechanism and provides a dynamical substrate for the propagation of microscopic fluctuations to the macroscopic scale (whole-cell) as a spark-induced-spark mediated Ca^2+^ wave (Figure 1Ai-ii). Due to the significant impact of the Ca^2+^ concentration in the junctional SR facilitating the feedback underlying Ca^2+^ sparks and their propagation, cardiac cells typically exhibit a minimum SR-Ca^2+^ load threshold above which whole-cell SCRE occur; luminal sensitisation of the RyRs [14] can also promote this highly SR-Ca^2+^ load dependence of SCRE.

Spontaneous Ca^2+^ release events are a potentially pro-arrhythmic cellular phenomenon: wholecell Ca^2+^ release can activate NCX which results in a transient inward current, depolarising the cell membrane potential as a delayed-after-depolarisation (DAD) or full triggered AP (TA; Figure 1Aii), which in tissue may present as a focal excitation. Simulating this behaviour therefore requires computational models which explicitly account for the underlying stochastic spatiotemporal dynamics (Figure 1Bi-ii), as have been recently developed by multiple groups from the super-resolution single-dyad to the whole-cell scale [14–33].

Despite the recent successes of these models to dissect cellular mechanisms of potentially proarrhythmic phenomena such as SCRE and Ca^2+^ transient alternans, only a few studies have attempted to dissect the tissue-scale considerations for the manifestation of these phenomena as arrhythmia, for example investigating: the minimum tissue substrate for the emergence of focal excitations resulting from non-stochastic EADs and DADs [34]; the emergence of focal excitation from stochastic SCRE [25,27,35–37] and its potential interaction with extracellular matrix remodelling [38]; SCRE as a mechanism for both triggered activity and conduction block [39]; and the potential complex considerations for pharmacological action on both triggers and substrate [27]. However, despite these important works, the role of SCRE in rapid arrhythmias at the organ scale is still to be fully explored, for example in relation to the direct translation of single cell data and modelling studies to the tissue-scale, and the multi-scale interaction of SCRE with re-entrant excitation.

Exploring the potential interactions between re-entry and SCRE using computational modelling presents a significant challenge due to the contrasting requirements of simulating each phenomenon: re-entry requires a minimum tissue size and therefore necessitates the use of efficient cell models, typically using the non-spatial Hodgkin-Huxley approach; 3D cell models capable of reproducing stochastic SCRE at dyad and whole-cell scale are computationally expensive to solve and unsuitable for simulation of the thousands or millions of coupled cells comprising tissue models appropriate for studying re-entry. Mean-field approximations can be applied to describe dyads as single structures [33] and distribute them throughout the free diffusion volume of a cell (Figure 1Bi, [26]), and subsequently to compartmentalise the intracellular and SR volumes surrounding a dyad as a Ca^2+^ release unit (CRU), arranged in an idealised cellular grid (Figure 1Bii, [16]). However, even these idealised cell models present computational requirements which are intractable for practical tissue-scale simulations with current computing power. The final level of model reduction from idealised spatial cell models to point-source Hodgkin-Huxley cell and resulting tissue models (Figure 1Biii) presents the largest challenge [11].

In this study, an approach was developed to overcome this challenge and dynamically model stochastic SCRE in non-spatial cellular and resulting tissue models, based on an extension of previous work [25]. A hierarchy of cell models was developed which contained both spatial and non-spatial equivalents, with analytical waveform functions used to reproduce observed dynamics of SCRE in the non-spatial cell models. The dynamics and variability of SCRE occurring in multiple conditions provided validation of the accurate reproduction of these stochastic dynamics at significantly reduced computational cost. Finally, the potential of the approach was demonstrated by assessing the vulnerability to focal excitation under different conditions.

**Figure 1:**
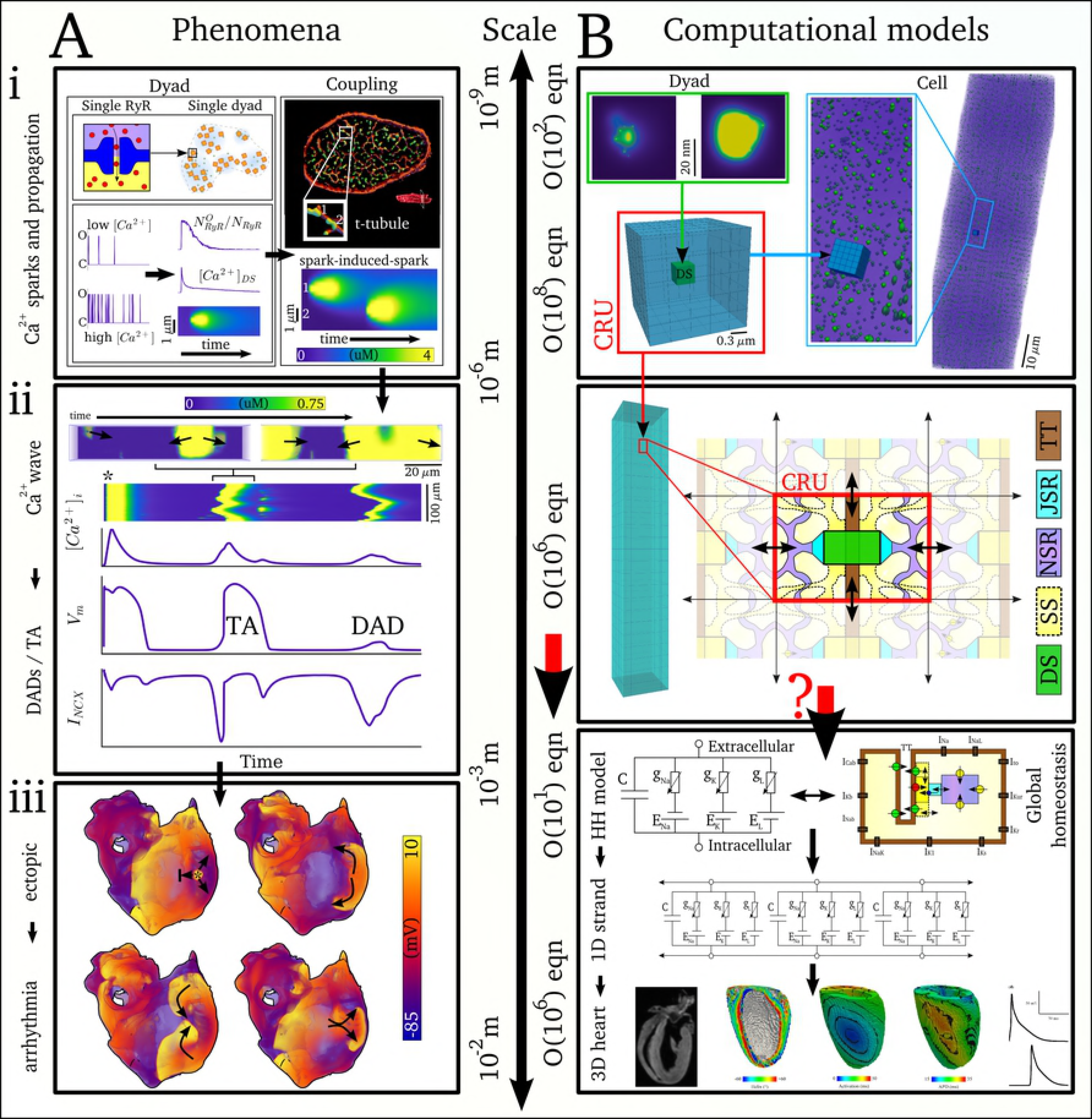
Multi-scale phenomena and computational modelling of cardiac calcium handling and electrophysiology. A Intracellular Ca^2+^ handling system and SCRE phenomena at different scales. A(i) single dyad (left panels): illustration of a single ryanodine receptor (upper left) and their distribution in a dyad (upper right); single channel open probability at low and high intracellular Ca^2+^ (lower left) and whole-dyad Ca^2+^ spark (lower right); panels on the right illustrate the T-tubule system in a cell cross-section (upper), the distribution of dyads (green) in the cross section and along a T-tubule (inset), and the propagation of a Ca^2+^ spark between two dyads along a T-tubule (lower). Unpublished structure data provided by Dr. Izzy Jayasinghe, University of Leeds; Ca^2+^ spark data from unpublished simulations. A(ii) Illustration of an intracellular Ca^2+^ wave and its effects on cellular excitation, showing two temporal snapshots of spatial Ca^2+^ concentration in a 3D volume (upper), a longitudinal linescan through the cell, and whole-cell voltage, Ca^2+^ and *I*_NCX_ associated with a stimulated excitation and two Ca^2+^ waves, the first initiating triggered activity (TA) and the second resulting only in a DAD. Simulation data from the present study used for illustration. (iii) Four temporal snapshots of excitation at the organ scale, demonstrating the relationship between tissue-level triggered activity (upper left) and the onset of re-entrant excitation through conduction block (remaining panels). Simulation data from Colman et al. 2013 [40]. B Computational modelling of cardiac electrophysiology at the different scales. B(i) Spatial model of a single dyad (upper left, unpublished data), with the remainder of the panel showing a structurally detailed model of a 3D cell in which dyads are modelled as single structures distributed through a free Ca^2+^ diffusion volume (data from Colman et al. 2017 [26]). B(ii) a coarse-grained 3D cell model in which the dyad and its surrounding intracellular spaces are considered as a single structure, the Ca^2+^ release unit (CRU). B(iii) Structure of a non-spatial, Hodgkin-Huxley type model illustrating: the electric circuit model of voltage dynamics (upper left) and non-spatial Ca^2+^ handling schematic (upper right); their coupling in 1D fibre strands (middle); and a reconstruction of cardiac anatomy for tissue simulations (lower panels data reproduced from Benson et al. 2011 [41]).

## Methods

This section first outlines the computational framework and the hierarchy of cell and tissue models of which it comprises. Then, analysis of stochastic SCRE observed in the spatial single cell models is discussed in relation to the development of analytical waveform functions which approximate SCRE in the non-spatial cell and multi-dimensional tissue models. Multiple implementations are presented in which the parameters are set by either: (i) directly controlled user inputs (“Direct Control” model); (ii) dynamically determined distributions to reproduce the behaviour of the spatial cell model under multiple conditions (“Dynamic Fit” model); or (iii) user controllable dynamically determined distributions (“General Dynamic” model).

### The computational framework

The developed framework consists of a hierarchy of computational models (Figure 2):

1. The microscopic or 3D spatio-temporal cell model is the baseline model for the framework. It accounts for spatially distributed dyads, Ca^2+^ diffusion within the cell, and stochastic state transitions in the LTCC and RyR models (Figure 2Ai, B). This model was used to study SCRE at the cellular scale.
2. The deterministic, 0D or non-spatial cell model is derived from the microscopic model. It does not contain a distributed Ca^2+^ handling structure or account for stochastic state transitions (Figure 2Aii, B). This manuscript focuses on the derivation and introduction of analytical Spontaneous Release Functions (SRF) into 0D models to reproduce behaviour from the 3D model.
3. The Tissue model refers to coupled 0D cell models in either idealised 2D sheets or 3D reconstructions of cardiac anatomy (Figure 2C).

**Figure 2:**
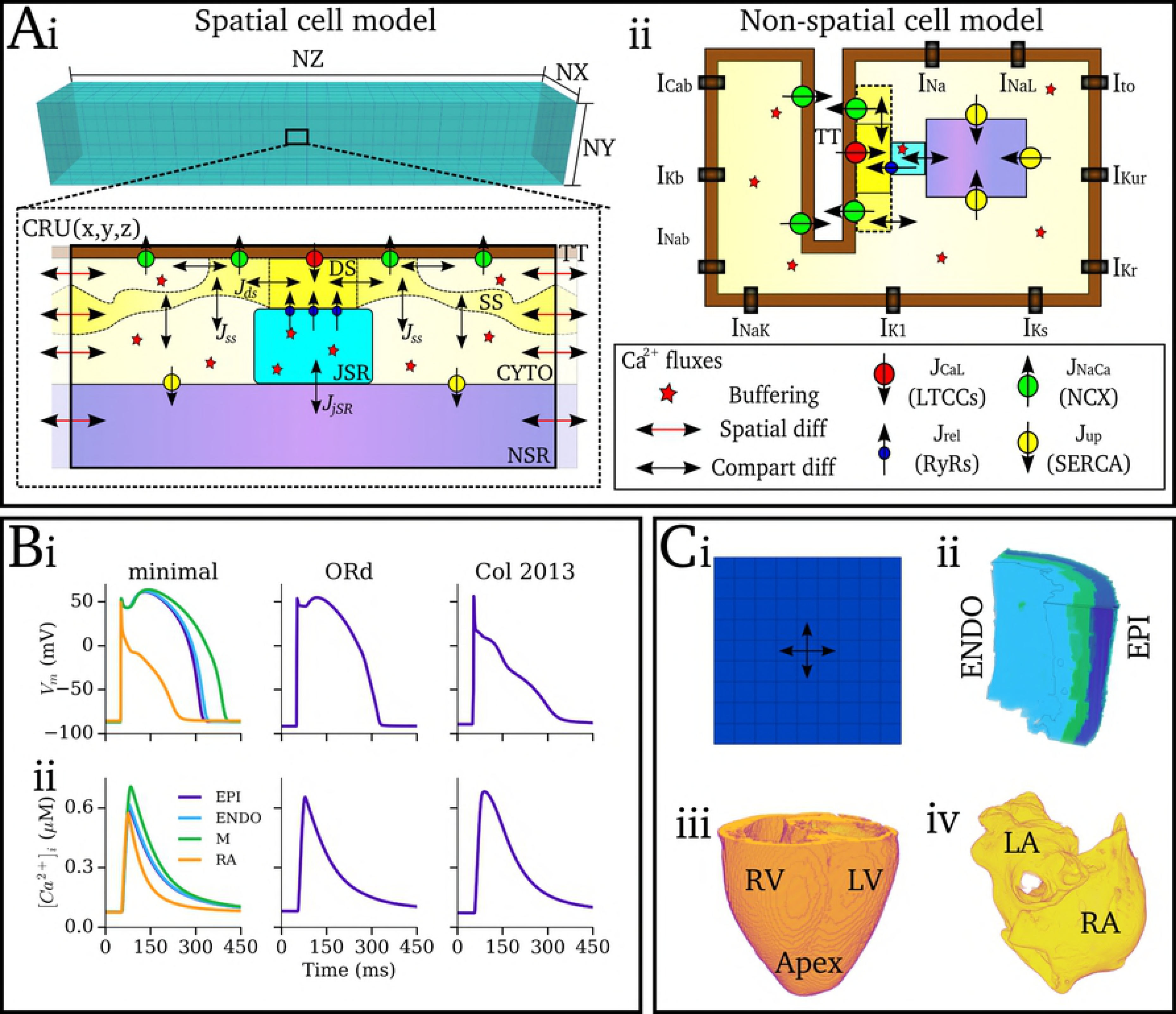
Components of the multi-scale computational framework. A Schematic of single cell Ca^2+^ handling and ion current models. (i) 3D, microscopic Ca^2+^ handling model, illustrating the 3D grid of calcium release units (CRUs; upper panel) and the compartments and Ca^2+^ fluxes within a single CRU (lower panel). Labelled are the dyadic cleft space (DS), sub-space (SS), bulk cytosolic space (CYTO), network and junctional SR spaces (NSR, JSR), and a T-tubule (TT); fluxes through the LTCCs (*J*_CaL_), RyRs (*J*_rel_), NCX (*J*_NaCa_) and SERCA (*J*_up_) are illustrated according to the key; double-headed black arrows indicate transfer between compartments; double-headed red arrows indicate diffusion between neighbouring CRUs. (ii) 0D, non-spatial cell model, illustrating the same fluxes as in (i) but without inter-CRU diffusion. Global ion currents are illustrated along the membrane (which apply to both models). B Whole-cell voltage (i) and calcium transient (ii) of the different ion-current models used in the present study, showing the minimal model (left), O’Hara et al., human ventricular model [42] (middle) and Colman et al., human atrial model [40] (right). C Tissue models, showing schematic of a 2D sheet model (i), and the 3D anatomical reconstructions: (ii) human ventricular wedge [43]; (iii) canine whole ventricle [44]; and (iv) human atria [40,45–47].

### The microscopic, 3D spatio-temporal Ca^2+^ handling model

The microscopic model, a simplification of a previously presented and structurally detailed model [26], consists of an idealised 3D intracellular Ca^2+^ system coupled to a point-source V_m_ and ioncurrent model (Figure 2Ai,B). The intracellular model consists of a 3D grid of CRUs, each containing five compartments (three intracellular compartments comprising the bulk-cytoplasm, sub-space, and restricted dyadic cleft; the network and junctional SR spaces; Figure 2Ai). The bulk cytosolic, sub-space and network SR compartments are coupled to neighbouring CRUs. Each dyad contains ~10 LTCCs and ~100 RyRs (varies between models and can be heterogeneous within a cell), described by stochastic differential equations [26]. Full details and model equations are provided in the Supplementary Material S1 Text (Model Description). Fundamental equations are given below:

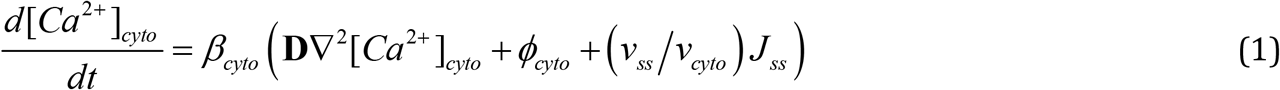

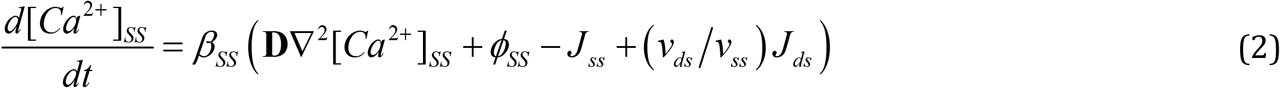

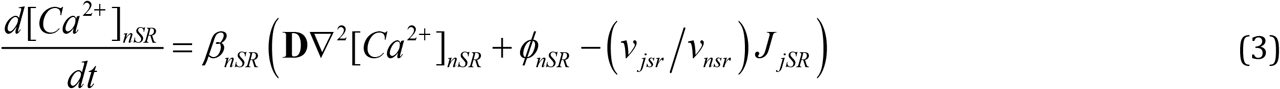

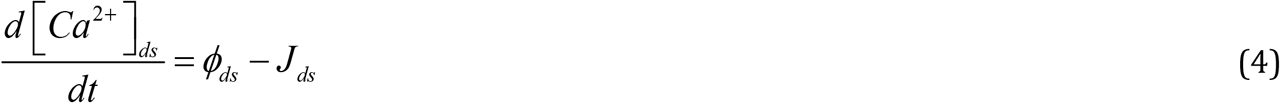

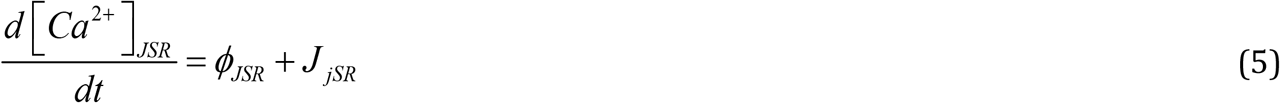

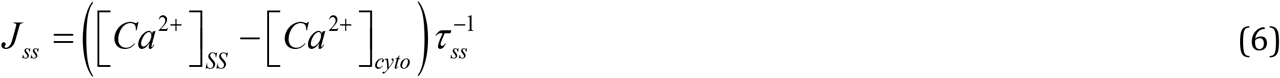

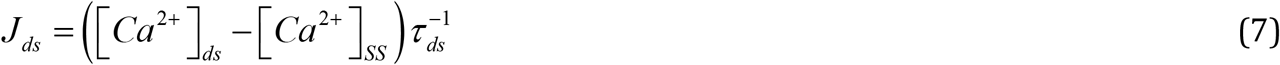

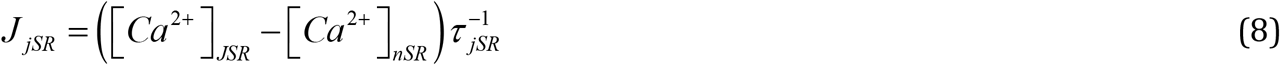

Where *Φ*_*x*_ is a general reaction term comprising the relevant channel fluxes, *β*_*x*_ is the instantaneous buffering term [48], *v*_*x*_ is the volume of the compartment in the subscript, *τ*_x_ is the time constant of inter-compartmental diffusion, *∇*^2^ is the spatial Laplacian operator in 3D and ***D*** is the diffusion coefficient. Full reaction terms are given in the Supplementary Material S1 Text (Model Description). Most relevant for this study is the reaction for the dyadic cleft, *Φ*_*ds*_, which contains intracellular Ca^2+^ release, *J*_rel_. For a single dyad *n, J*_rel_ is described by:

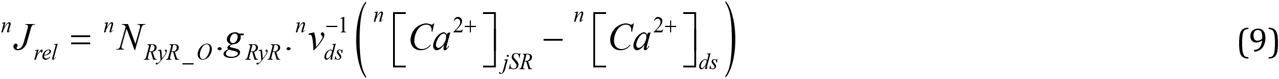

Where *g*_RyR_ is the maximal flux factor for Ca^2+^ release, *v*_ds_ is the volume of the dyadic cleft space, and ^n^*N*_RyR_O_ is the summed number of open RyR channels in dyad *n*, determined through the stochastic Monte-Carlo method. *N*_RyR_O_ is therefore the primary variable which describes the dynamics of SCRE.

### The Deterministic, 0D model

The deterministic model structure is identical to a single CRU of the microscopic model and of the same form as the majority of contemporary cardiac cell models based on the Hogkin-Huxley approach (Figure 2Aii). Both the RyR and LTCC models are described by ordinary differential equations and solved through the deterministic forward-Euler method. Modifications to the RyR model were required due to the poor recapitulation of whole cell CICR using deterministic approaches see Supplementary Material S1 Text (Model Description) for details.

### Action potential and tissue models

For the purpose of demonstrating the general potential of the developed framework for integration with contemporary cell models, multiple ionic models were integrated with the Ca^2+^ handling system to describe the non-Ca^2+^ dependent membrane currents (Figure 2B): the model was integrated with simplified versions of the O’Hara et al., 2011 human ventricular AP model [42] and Colman et al., 2013 human atrial AP model [40,49]. Furthermore, a hybrid-minimal model (comprising of a minimal setup suitable for coupling with physiological Ca^2+^ currents) was developed which describes human AP morphology in the three transmural cell types of the ventricles as well as the atrial myocardium.

Idealised 2D sheet models consist of a 2D array of coupled cells in isotropic medium (Figure 2Ci), including both homogeneous sheets of either the ventricular epicardial layer or the right atrial wall, and a model of the transmural heterogeneity in the ventricular wall (simple ratio of 1:1:1 ENDO:M:EPI cells). 3D models consist of a wedge reconstruction of the human ventricular wall (Figure 2Cii, [43]), a reconstruction of the whole canine ventricle [44] (Figure 2Ciii), and a reconstruction of the whole human atria [40,45–47] (Figure 2Civ).

### Pro-arrhythmic conditions and analysis protocols

To induce prominent full-cell release events through different cellular conditions, representative (but non-specific) models were included for isoprenaline (ISO, sympathetic response which enhances CICR) and two types of pro-SCRE general disease remodelling mimicking features observed in conditions such as AF and HF (e.g., [9,50]): (i) SERCA was up-regulated and NCX was down-regulated (R_SERCA/NCX_); (ii) the SR-Ca^2+^ threshold for release was lowered through increased inter-CRU coupling (R_CRU-CRU_). R_SERCA/NCX_ also included remodelling of the ion currents to provide a different AP environment coupled to the SCRE, including a reduction of *I*_K1_. Details are provided in the Supplementary Material S1 Text (Model Description). These models were combined with a rapid pacing protocol to promote SR-Ca^2+^ loading and the emergence of whole-cell SCRE: The models were paced to stable state under varying basic cycle lengths (BCL = 200-600 ms); the state variables were saved at the stable values and used as initial conditions to efficiently run any number of simulations of a short pacing period followed by a quiescent (non-applied pacing) period, within which the statistics of SCRE can be analysed.

Furthermore, a Ca^2+^-clamp ladder protocol (Figure 3) was implemented to both derive and validate the SRF in the 0D model: At each step, the intracellular and SR-Ca^2+^ concentrations were held at specified values for 2 s, with the SR-Ca^2+^ concentration increasing for each step in succession (Figure 3Aii). When intracellular release occurs, the Ca^2+^ concentrations were allowed to dynamically evolve. The membrane potential was allowed to evolve during this protocol, but the conductances of *I*_Na_ and *I*_CaL_ were set to zero to prevent excitation and resulting interruption of the SCRE by CICR. This protocol illustrates the variety of SCRE and their underlying spatiotemporal dynamics which must be captured in the 0D model (Figure 3B; Supplementary Material S1 Video [Ca^2+^ clamp illustration]).

**Figure 3:**
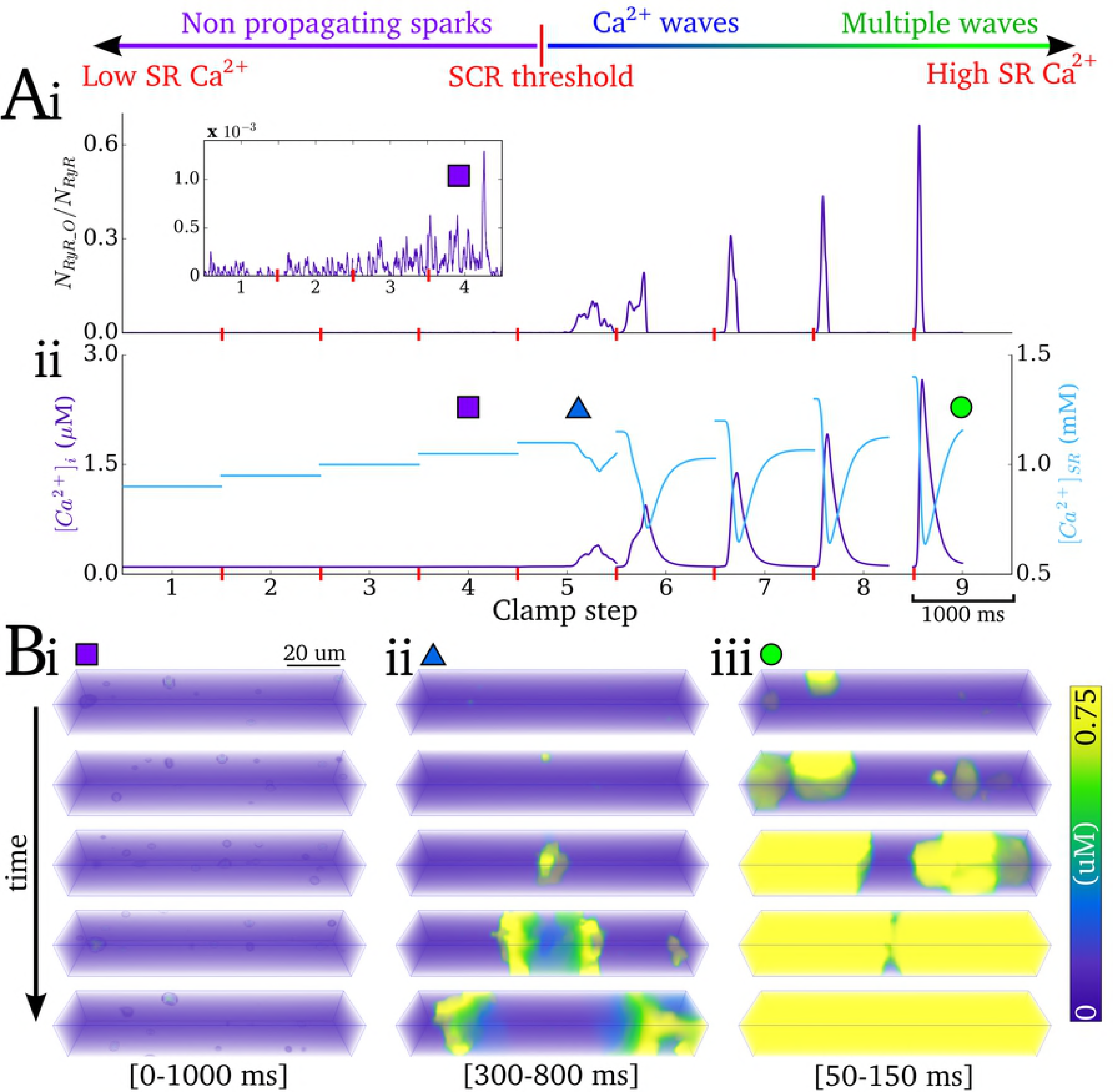
Illustration of Ca^2+^ clamp protocol. A - Ca^2+^ clamp protocol illustrated for 9 steps of SR-Ca^2+^, showing traces for: (i) proportion open RyR; (ii) intracellular(purple) and SR(blue) Ca^2+^ concentration. B Snapshots of the spatio-temporal Ca^2+^ dynamics at different SR-Ca^2+^ concentrations, showing: (i) non-propagating sparks; (ii) slow Ca^2+^ wave; (iii) multiple and rapid Ca^2+^ waves; the time range for the snapshots is shown in the square brackets.

### Derivation of the Spontaneous Release Functions (SRF)

The phenomenological approach involved the development of SRF which describe RyR dynamics associated with SCRE observed in the 3D cell model. These functions are waveforms associated with the range of open RyR morphology (Figure 4A).

**Figure 4:**
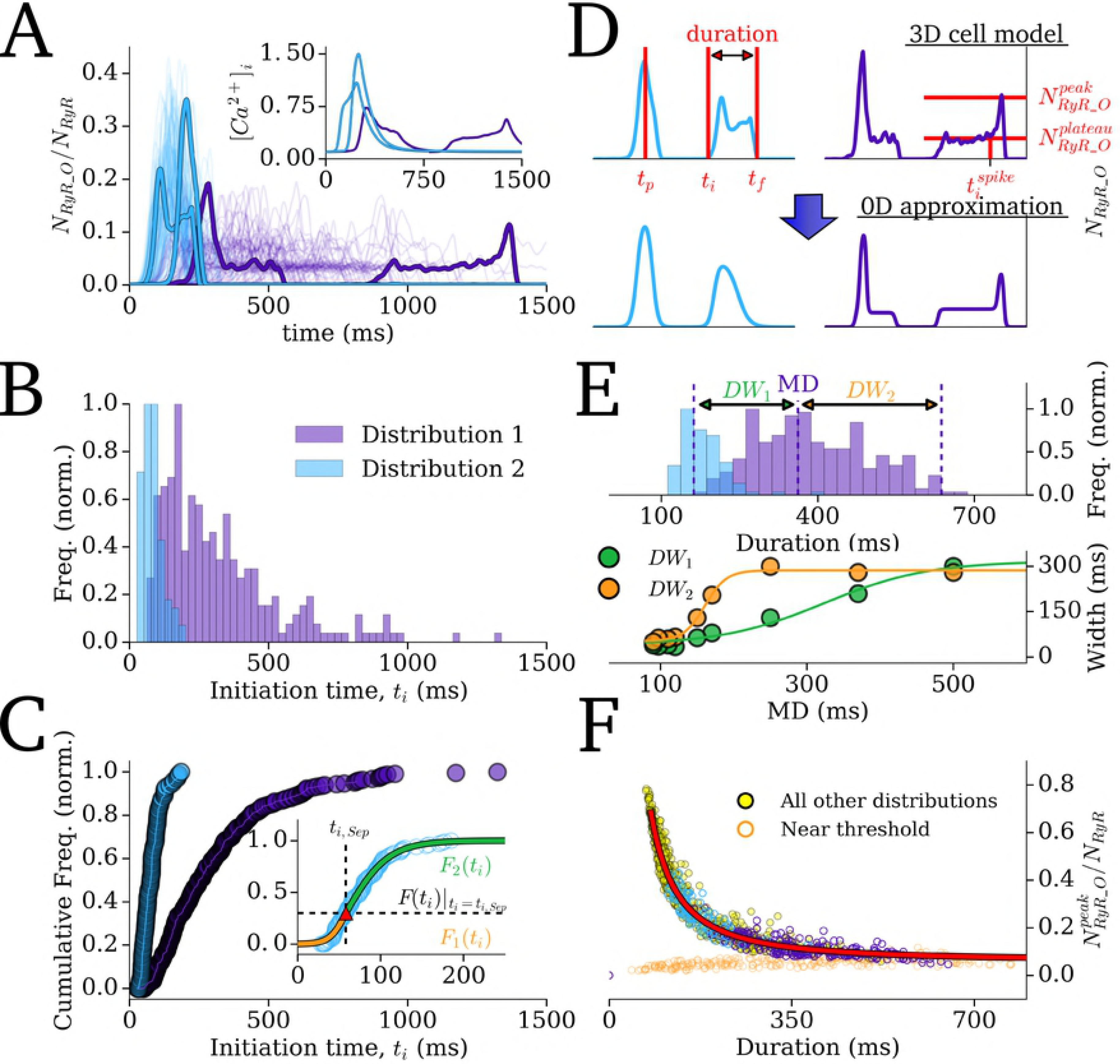
Derivation of the Spontaneous Release Functions. A - Traces of open RyR associated with 100 SCREs at a low (purple) and high (blue) SR-Ca^2+^ in the 3D cell model. B - Histogram describing the initiation time for SCRE and C - the correlating cumulative frequency plots, associated with 250 simulations of each condition. Inset - example of fitting the cumulative frequency with two sigmoidal functions (*F*_1_(t_i_) and *F*_2_(t_i_), orange and green), separated at a specific point (*t*_i_Sep_, *F*(_ti_)|_ti=ti_Sep_, red triangular marker). D - Examples of two types of waveform (upper panel) and the SRF which approximate them (lower panel). Labelled are the parameters which fully describe the waveforms: the initiation time (*t*_i_), peak time (*t*_p_), final time (*t*_f_), initiation time of spike during plateau (*t*_i_^spike^), peak open RyR (*N*_RyR_O_^peak^), plateau open RyR (*N*_RyR_O_^plateau^). E Histogram illustrating the distribution of RyR waveform duration for the two conditions, with the median duration (*MD*) and width of distribution either side of the median (*DW*_1_, *DW*_2_) labelled for Distribution 1 (upper panel); relationship between the distribution widths and median (lower panel, points data; lines fit by equations (18,19)). F Correlation between peak of open RyR and the duration, shown for the two conditions featured in the Figure (purple, blue) and all other simulations (yellow). Low amplitude SCRE occurring near the threshold SR-Ca^2+^ are shown in orange. Fit by equation (20) is shown by the red line.

#### RyR waveforms

Results from the Ca^2+^ clamp ladder protocol across all SR-Ca^2+^ values were used to derive the analytical formulations of the SRF describing the variability in spontaneous release *N*_RyR_O_ waveforms. Comparison of the SCRE waveforms from 250 simulations at low and high SR-Ca^2+^ illustrates the range of waveforms observed (Figure 4A) and the corresponding probability density function and cumulative frequency of initiation times, *t*_i_ (Figure 4B-C).

The *N*_RyR_O_ waveforms can be grouped into two primary types: spike-like associated with short, large-amplitude release, and plateau-like associated with long, small-amplitude release (Figure 4D). For the spike-like morphology, the waveform can be well approximated with the simple function:

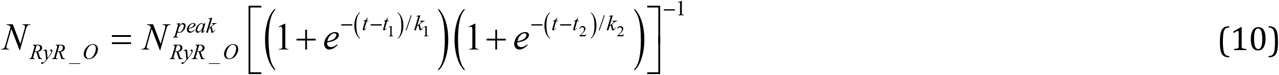

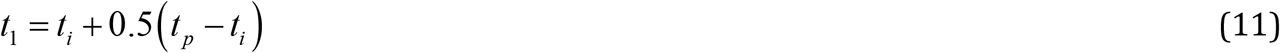

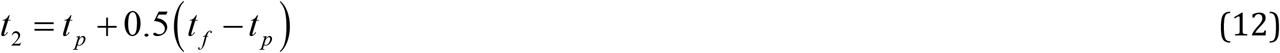

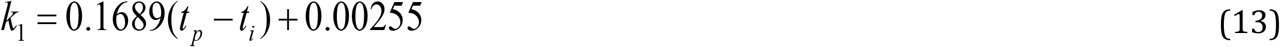

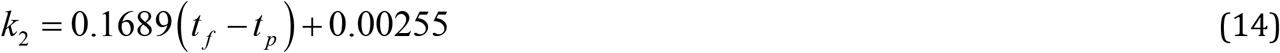

where *t*_*i*_ is the initiation time of the SCRE, *t*_f_ is the end time (duration, *λ*, thus = *t*_f_-*t*_i_), *t*_p_ is the time of the peak of the waveform and *N*_RyR_O_^peak^ is the peak of open proportion RyR (Figure 4D). The function for the plateau-like waveform (corresponding to durations longer than 300 ms) is derived from the same parameters:

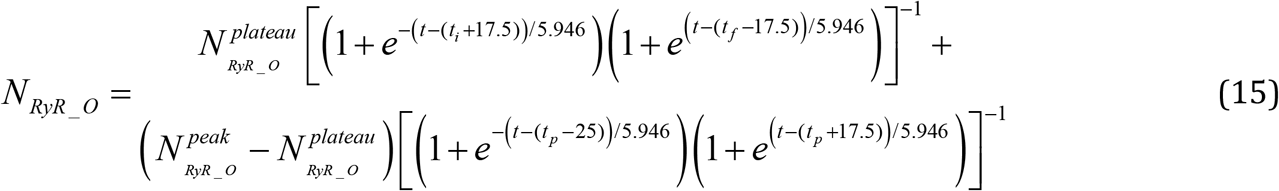

Where *N*_RyR_O_^plateau^ is the amplitude of the plateau (Figure 4D). This equation assumes the same form for the spike occurring within the plateau, with its upstroke time being 50 ms and its decay time 35ms; *t*_i_^spike^ (Figure 4D) therefore corresponds to *t*_p_-50 (and its half maximal activation time *t*_p_-25).

The waveform is therefore completely described by four-five parameters: (1) initiation time, *t*_i_; (2) duration (*λ = t*_f_−*t*_i_); (3) peak time, *t*_p_; and (4-5) amplitude (*N*_RyR_O_^peak^; *N*_RyR_O_^plateau^). In order to maintain physiological waveforms and randomly sample the parameter values from appropriate distributions, the nature of stochastic variation of these four parameters is discussed below.

#### Parameter distributions

1. *t*_i_: The probability density functions for the initiation time associated with each SR-Ca^2+^ value do not demonstrate a normal distribution, but rather a skewed distribution (Figure 4B). The cumulative frequency (Figure 4C) is well approximated by the use of two simple sigmoidal functions (Figure 4C-inset), maintaining the desire for restraint in the number of parameters and allowing simple and intuitive controllability:

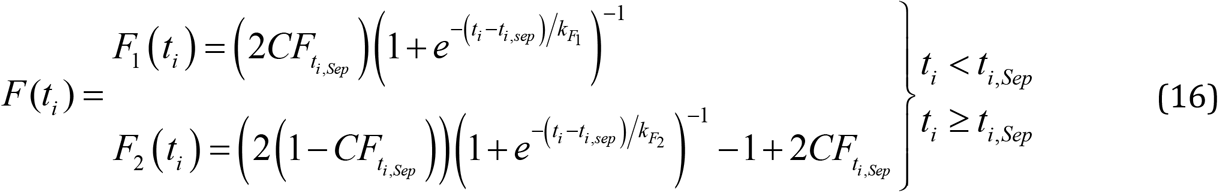 The distribution for *t*_i_ is therefore determined by four parameters: the initiation time corresponding to the point where the functions are separated (*t*_*i,Sep*_); the cumulative frequency at this point (*CF*_ti,Sep_= *F*(*t*_i_)|_ti=ti,Sep_), and the gradient parameter of each function (*k*_F1_, *k*_F2_ corresponding to the width of the distribution either side of *t*_i,Sep_; Figure 4C-inset.)
2. *λ*: The distributions for the duration are also non-normal, and well approximated by two sigmoidal functions describing the cumulative frequency for half of the data either side of the median duration (*MD*; Figure 4E):

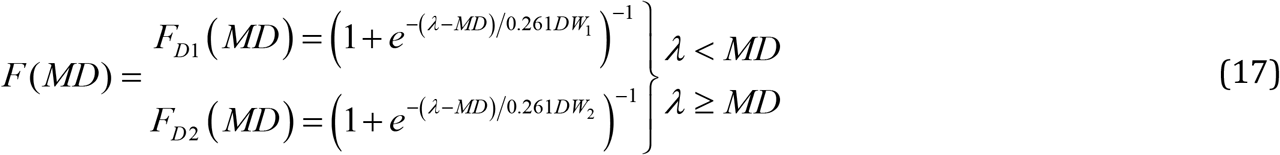

Where the widths (*DW*_1_, *DW*_2_, in ms) are a function of the *MD* (Figure 4E), given by:

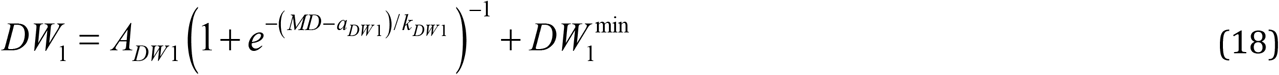

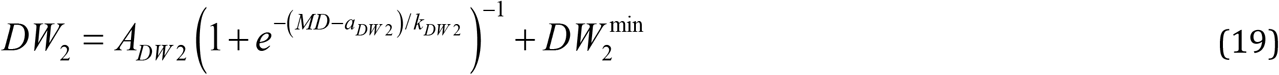

Default parameters are given in Table 1. The duration distribution is therefore completely described by the median, *MD*. Note that the widths (*DW*_1_, *DW*_2_) could also be specified directly for complete control over the variability in duration.
3. *t*_p_: The timing of the peak varies approximately evenly within the duration of the wavefrom, occurring between 25 ms after the initiation (*t*_i_) and 52 ms before the final time (*t*_f_).
4. *N*_RyR_O_^peak^; *N*_RyR_O_^plateau^: The amplitude correlates strongly with duration, λ (Figure 4F):

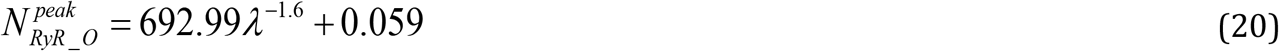

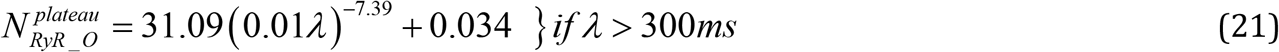

With small variance (± 0.05; ± 0.25*N*_RyR_O_^plateau^). Note that it would not be appropriate to define the amplitude independently from the duration, due to the correlation between these two parameters corresponding to the total amount of Ca^2+^ released.

#### The Direct Control SRF model

With this setup, therefore, all parameters of the waveform are derived from two primary waveform properties: the initiation time, *t*_i_, and the duration, *λ* (which also determine the peak time and amplitude); the distributions describing the variability of these properties are entirely described by 5-7 parameters (*t*_i_ = *f*(*t*_i_sep_, *CF*_ti_sep_, *k*_F1_, *k*_F2_); *λ* = *f*(*MD, DW*_1_, *DW*_2_) where *DW*_1_, *DW*_2_ = *f*(*MD*) or specified). For the simplest implementation of SCRE in 0D models, the user Direct Control model, SRF variability and morphology can therefore be described by simply defining these 5-7 parameters (as well as a probability of SCRE) in order to reproduce single conditions (for example, fit to a single dataset). The next section describes an approach to derive these parameters dynamically based on relevant Ca^2+^ handling environment variables.

#### The Dynamic Fit SRF model

The Dynamic Fit SRF model was derived through correlation of the parameters defining the *t*_i_ and *λ* distributions with the primary environment variable controlling SCRE: the SR-Ca^2+^ concentration (Figure 5). Relation to this single variable was chosen for practicality and simplicity of the resulting equations, which is in particularly valuable for reproducing variable Ca^2+^ handling system states.

**Figure 5:**
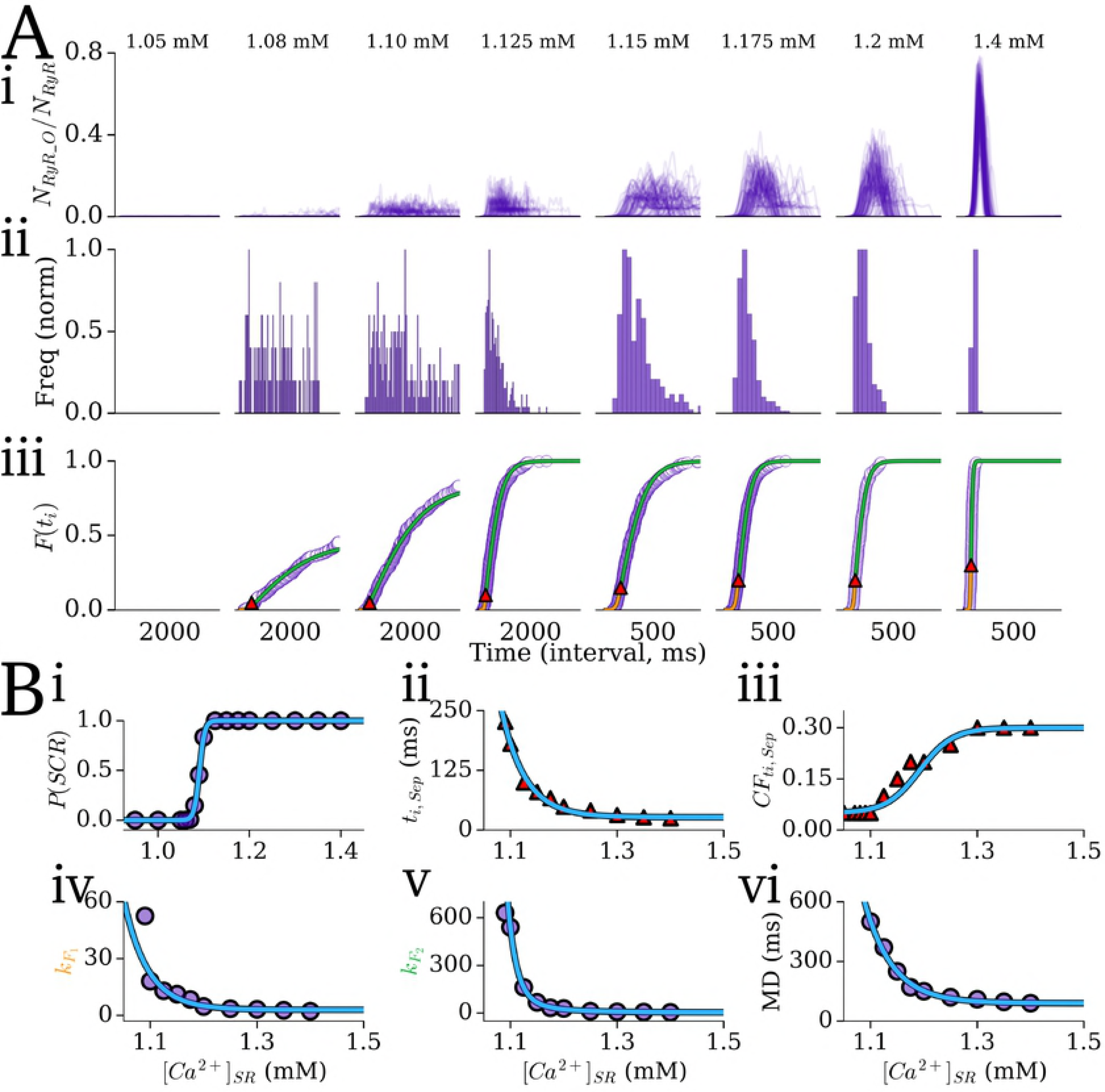
Derivation of the Dynamic SRF parameters in the control model. A - RyR waveforms from 100 simulations of the Ca^2+^ clamp protocol (i) over eight steps of SR-Ca^2+^ (panel titles), and corresponding initiation time distributions (ii) and cumulative frequency plots (iii) from 250 simulations at each SR-Ca^2+^. The values given on the *x-axis* refer to the total time interval of the plot, not absolute values. The two fitting functions (*F*_1_(t_i_) and *F*_2_(t_i_), orange and green; see Figure 4) and their separation point (red triangular marker) are shown. B - Summary data (purple dots, red triangles) and the fit from the relevant functions (blue line) against SR-Ca^2+^ for: (i) probability of whole-cell SCRE; (ii) initiation time corresponding to the separation point, *t*_i,Sep_; (iii) the normalised cumulative frequency at this point, *CF*_ti_Sep_= *F*(_ti_)|_ti=ti_Sep_; (iv) the *k* parameter for *F*_1_(t_i_) and (v) for *F*_2_(t_i_); (vi) the median duration, *MD*.

The probability that an SCRE occurs at a given SR-Ca^2+^ is well-approximated by a sigmoidal function (Figure 5Bi):

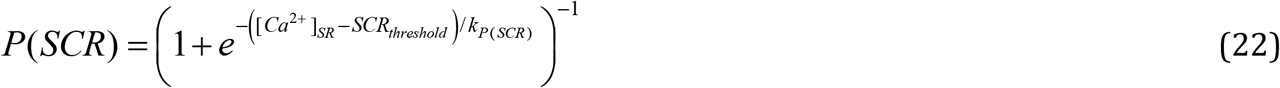

The SR-Ca^2+^ dependence of *t*_i_sep_ and *CF*_ti_Sep_ can be approximated by the following functions (Figure 5Bii,iii):

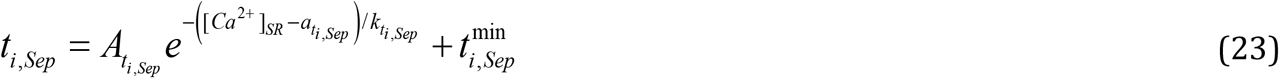

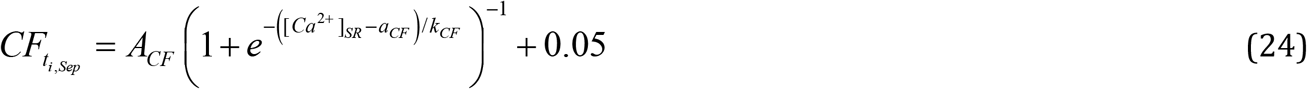

The gradient parameters for *F*_1_(t_i_) and *F*_2_(t_i_), *k*_F1_ and *k*_F2_, are then approximated by the following functions (Figure 5Biv,v):

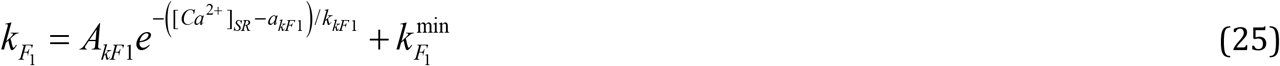

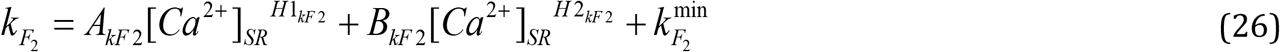

And finally, the MD also correlates well with the SR-Ca^2+^:

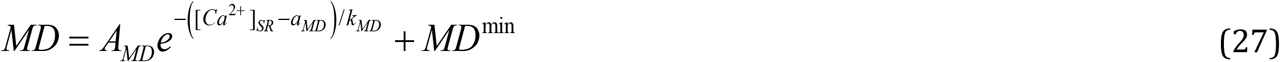

Parameters corresponding to the control model are given in Supplementary Material S1 Text (Model Description). Use of these simple form functions easily allows re-fitting to data from different conditions and thus the derivation of these SRF parameters for the two remodelling conditions, both of which directly affect the sub-cellular dynamics of SCRE (Figure 6; Table 1).

**Figure 6:**
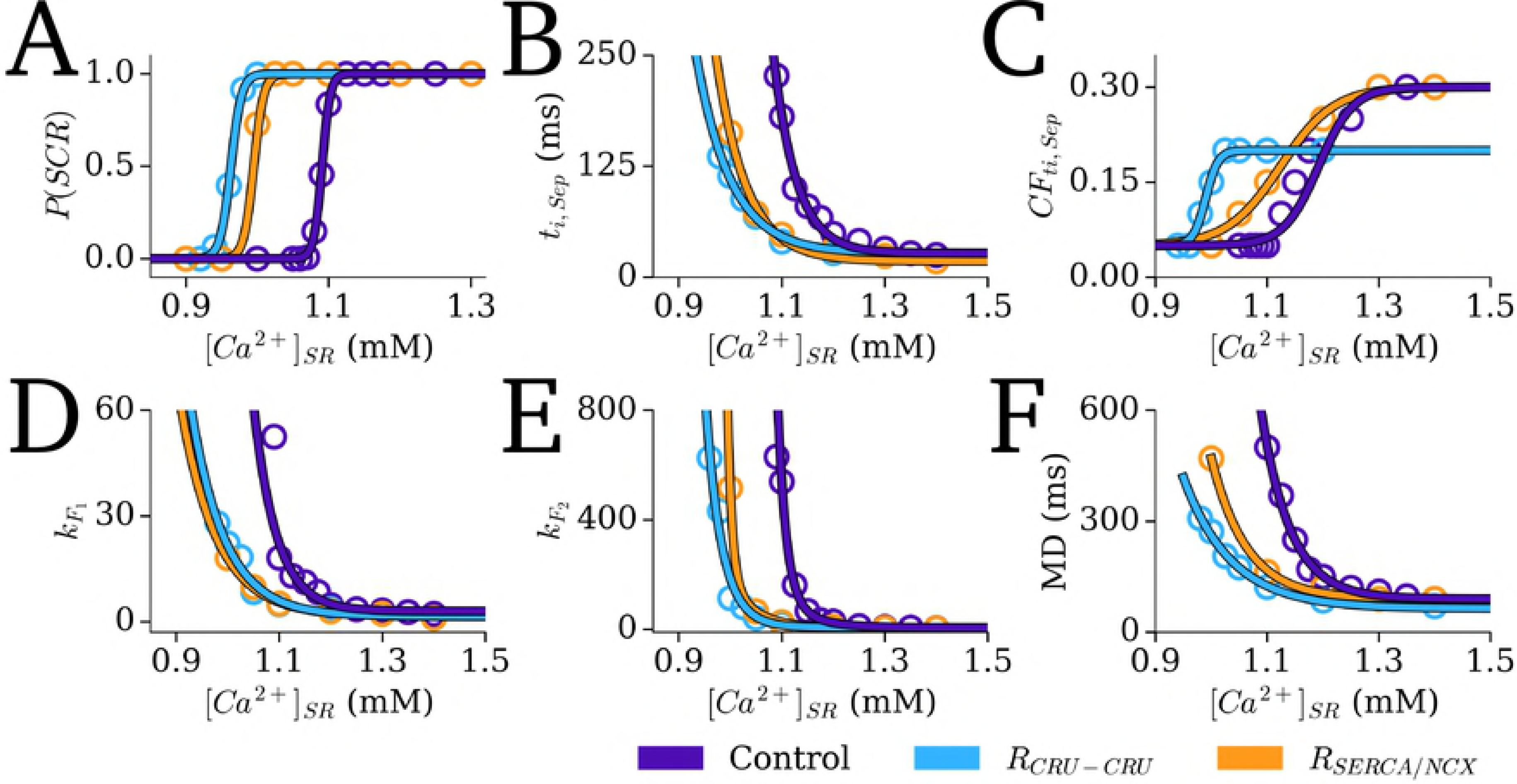
Incorporating different Ca^2+^ handling conditions into the Dynamic SRF. Summary data (points) and the fit from the relevant functions (lines) for the three Ca^2+^ handling conditions (purple - control; blue - CRU-CRU coupling enhancement; orange - SERCA upregulation and NCX downregulation model) against SR-Ca^2+^ for: A - probability of whole-cell SCRE; B - initiation time corresponding to the separation point, *t*_i_; C - the normalised cumulative frequency at this point, *CF*_ti,Sep_= *F*(_ti_)|_ti=ti,Sep_; D - the *k* parameter for *F*_1_(t_i_) and (v) for *F*_2_(t_i_); (vi) the median duration, *MD*.

**Table 1.** Spontaneous release function SR-Ca^2+^ dependence parameters.

| Symbol | Parameter | Value<br>control | Value<br>$R_{\text{SERCA/NCX}}$ | Value<br>$R_{\text{CRU-CRU}}$ |
| --- | --- | --- | --- | --- |
| $SCR_{\text{threshold}}$ | SR-$\text{Ca}^{2+}$ threshold for SCRE (mM) | 1.091 | 0.993 | 0.963 |
| $k_{P(\text{SCR})}$ | Gradient parameter for $P(\text{SCR})$ | 0.0058 | 0.007 | 0.0075 |
| $A_{t_{i,\text{Sep}}}$ | Coefficient for $t_{i,\text{Sep}}$ function | 37.78 | 27.62 | 65.047 |
| $a_{t_{i,\text{Sep}}}$ | $t_{i,\text{Sep}}$ exponent $[\text{Ca}^{2+}]_{\text{SR}}$ reference parameter | 1.157 | 1.096 | 1.017 |
| $k_{t_{i,\text{Sep}}}$ | $t_{i,\text{Sep}}$ exponent $[\text{Ca}^{2+}]_{\text{SR}}$ gradient parameter | 0.041 | 0.0581 | 0.0674 |
| $t_{i,\text{Sep}}^{\min}$ | Minimal value for $t_{i,\text{Sep}}$ | 31.83 | 18 | 27 |
| $A_{\text{CF}}$ | Coefficient for $\text{CF}_{t_{i,\text{Sep}}}$ function | 0.25 | 0.25 | 0.15 |
| $a_{\text{CF}}$ | $\text{CF}_{t_{i,\text{Sep}}}$ exponent $[\text{Ca}^{2+}]_{\text{SR}}$ reference parameter | 1.192 | 1.124 | 0.99 |
| $k_{\text{CF}}$ | $\text{CF}_{t_{i,\text{Sep}}}$ exponent $[\text{Ca}^{2+}]_{\text{SR}}$ gradient parameter | 0.0311 | 0.0481 | 0.0111 |
| $A_{k_{\text{F1}}}$ | Coefficient for $k_{\text{F1}}$ function | 8.619 | 3.955 | 10.93 |
| $a_{k_{\text{F1}}}$ | $k_{\text{F1}}$ exponent $[\text{Ca}^{2+}]_{\text{SR}}$ reference parameter | 1.136 | 1.1 | 1.04 |
| $k_{k_{\text{F1}}}$ | $k_{\text{F1}}$ exponent $[\text{Ca}^{2+}]_{\text{SR}}$ gradient parameter | 0.0449 | 0.0704 | 0.0659 |
| $k_{\text{F1}}^{\min}$ | Minimal value for $k_{\text{F1}}$ | 2.967 | 1.532 | 1.529 |
| $A_{k_{\text{F2}}}$ | 1<sup>st</sup> coefficient for $k_{\text{F2}}$ function | 583.6 | 173.361 | 181.12 |
| $B_{k_{\text{F2}}}$ | 2<sup>nd</sup> coefficient for $k_{\text{F2}}$ function | 295149 | 338.742 | 0 |
| $H1_{k_{\text{F2}}}$ | $[\text{Ca}^{2+}]_{\text{SR}}$ power for 1<sup>st</sup> term $k_{\text{F2}}$ function | -18.979 | -21.099 | -31.02 |
| $H2_{k_{\text{F2}}}$ | $[\text{Ca}^{2+}]_{\text{SR}}$ power for 2<sup>nd</sup> term $k_{\text{F2}}$ function | -68.306 | -118.771 | N/A |
| $k_{\text{F2}}^{\min}$ | Minimal value for $k_{\text{F2}}$ | 5 | 3.9 | 6.7 |
| $A_{\text{MD}}$ | Coefficient of median duration (MD) function | 208.57 | 801.86 | 136.86 |
| $a_{\text{MD}}$ | $\text{MD}$ exponent $[\text{Ca}^{2+}]_{\text{SR}}$ reference parameter | 1.136 | .953 | 1.0312 |
| $k_{\text{MD}}$ | $\text{MD}$ exponent $[\text{Ca}^{2+}]_{\text{SR}}$ gradient parameter | 0.0526 | 0.063 | 0.0854 |
| $\text{MD}^{\min}$ | Minimal value of MD | 90 | 90 | 64.12 |
| $A_{\text{DW1}}$ | Coefficient for $\text{DW}_1$ function | 276.72 | 273.53 | 155.61 |
| $a_{\text{DW1}}$ | $\text{DW}_1$ exponent MD reference parameter | 324.11 | 268.012 | 223.98 |
| $k_{\text{DW1}}$ | $\text{DW}_1$ exponent MD gradient parameter | 70.67 | 40.24 | 57.27 |
| $\text{DW}_1^{\min}$ | Minimal value of $\text{DW}_1$ | 40 | 48.26 | 40 |
| $A_{\text{DW2}}$ | Coefficient for $\text{DW}_2$ function | 233.26 | 369.44 | 109.18 |
| $a_{\text{DW2}}$ | $\text{DW}_2$ exponent MD reference parameter | 160.74 | 188.14 | 140.46 |
| $k_{\text{DW2}}$ | $\text{DW}_2$ exponent MD gradient parameter | 14.6 | 18.72 | 10.7 |
| $\text{DW}_2^{\min}$ | Minimal value of $\text{DW}_2$ | 53 | 60.56 | 70.422 |

### Implementation with the 0D cell model

These SRF were implemented within the 0D cell models using a simple algorithm (Figure 7): the input parameters (*P*(SCR), *t*_i,Sep_, *CF*_ti,Sep_, *k*_F1_, *k* _F2_, *MD*; see previous section: Parameter distributions) are defined at a certain time (see below) and then the waveform parameters (*t*_i_, and *λ*, which in turn define *t*_p_ and *N*_RyR_O_^peak^; see previous section: Parameter distributions) are randomly sampled from the associated distributions. When the SRF have been initiated (i.e., *t* > *t*_i_ and *N*_RyR_O_SRF is > 0), the *N*_RyR_O_ in the cell model is set to this value and the model thus evolves as if the equivalent SCRE was occurring in the 3D cell model. The inverse functions, giving the specific parameter value from a random number input and the distribution parameters, are as follows:

Initiation time, *t*_i_; inverse function of equation (16):

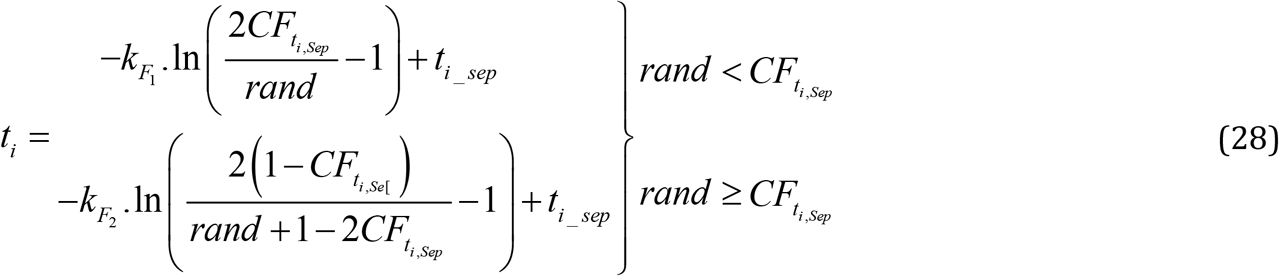

Duration, *λ*; inverse function of equation (17):

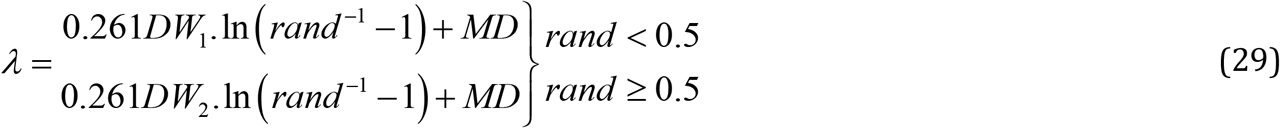

Peak time, *t*_p_, following a uniform distribution approximation:

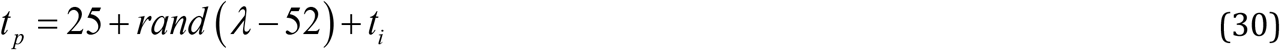

*N*_RyR_O_^peak^, *N*_RyR_O_^plateau^ are determined from *λ* directly from their definition functions (eqns 20-21) and ±*rand* × simple variance magnitude of 0.05 and 0.25*N*_RyR_O_^plateau^, respectively. For the simplest implementation of the SRF, the Direct Control model, this calculation is determined at the time of cellular excitation, setting *t*_i_ and *λ* based on the distributions defined by user input parameters (*P*(SCR), *t*_i,Sep_, *CF*_ti_sep_, *k*_F1_, *k*_F1_, *MD*) and five random numbers input into the corresponding inverse functions: first, a random number between 0 and 1 is generated and compared to the probability of release, *P(SCR)*; if *rand* < *P(SCR)*, then the input parameters are used to determine the inverse functions (equations 28-30) and 4 more random numbers determine the actual SRF waveform parameters; if *rand* > *P(SCR)*, then no SRF parameters are set. Note that the *t*_i_sep_ must be set relative to excitation time. The model will set SRF parameters based on these single distributions with every cellular excitation (note that the parameters may give an SCRE timing later than the next stimulated excitation, in which case it will be reset on the next excitation).

For the Dynamic Fit model, the calculation is performed multiple times, dynamically determined during the simulation: After an initial stimulated AP, when the RyR availability has recovered above a set threshold, the SR-Ca^2+^ is input to first define the probability of release, *P(SCR)*, from equation (22) and the same process is followed as described above, with the remaining input parameters defined from the SR-Ca^2+^ according to the appropriate functions [equations 23-27)]. If the SR-Ca^2+^ concentration changes more than by a predefined value (e.g., 0.01 mM) before the SCRE has been initiated, then the parameters are recalculated based on this new SR-Ca^2+^. SRF parameters are not calculated during excitation (i.e., when the RyR availability is low) but recalculated upon RyR recovery.

**Figure 7:**
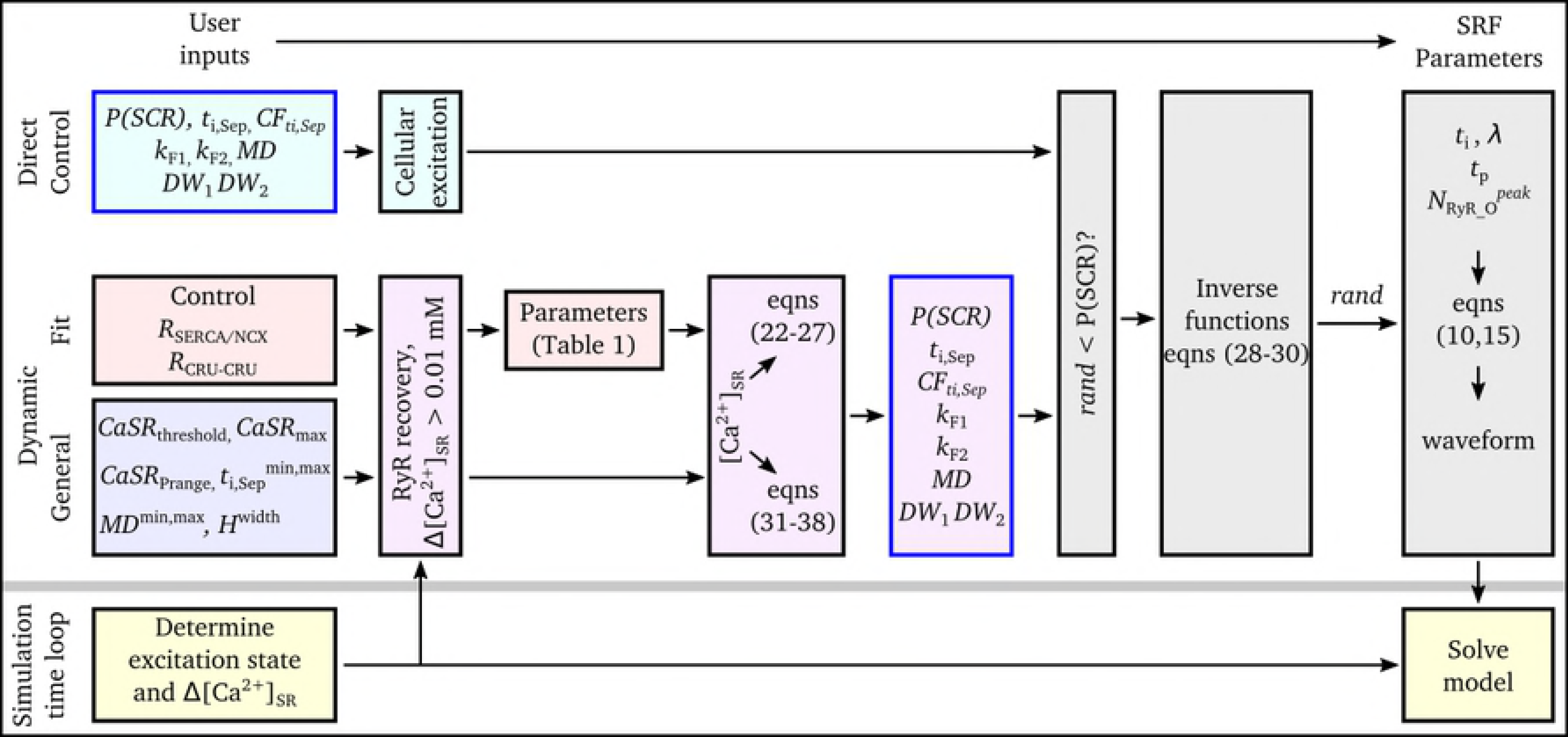
Implementation algorithm. Schematic of the algorithms used to integrate the SRF with the non-spatial cell models for the three implementations.

### The General Dynamic SRF model

Whereas the Dynamic Fit SRF model described above ensures congruence between the 3D and 0D models, a General Dynamic SRF model was also derived which provides full control over the distributions determining the dynamic behaviour. The approach therefore involved defining the functions which correlate SR-Ca^2+^ with the SRF input parameters (*P*(SCR), *t*_i_sep_, *CF*_ti,Sep_, *k*_F1_, *k*_F1_, *MD*), which define the inverse functions from which the actual waveform parameters are sampled (equations 28-30), based on intuitive controllable parameters (Figure 8): (i) - The threshold for SCRE (*CaSR*_threshold_); (ii) - The SR-Ca^2+^ range over which *P*(SCR) varies from 0 to 1 (*CaSR*_P_range_); (iii) - The maximal SR-Ca^2+^ above which SCRE distributions converge (*CaSR*_max_); (iv) - The minimum and maximum *t*_i,Sep_ and *MD* (*t*_i,Sep_^min^, *t*_i,Sep_^max^, *MD*^min^, *MD*^max^); (v) - The *t*_i_ and *λ* distribution widths at these extremes (*t*_i,width_^min^, *t*_i,width_^max^, *λ*_width_^min^, *λ*_width_^max^); And (vi) the non-linearity of width variance (*H*_width_); *CF*_ti,Sep_ was set to 0.4. The General Dynamic SRF parameters are therefore determined by the following equations:

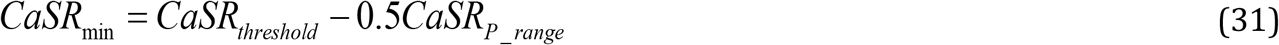

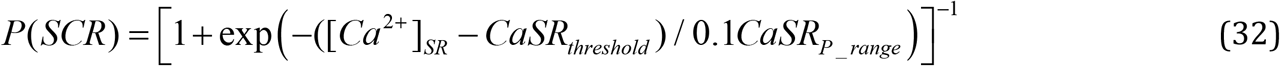

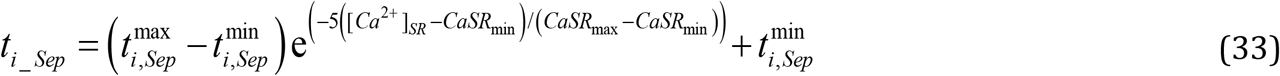

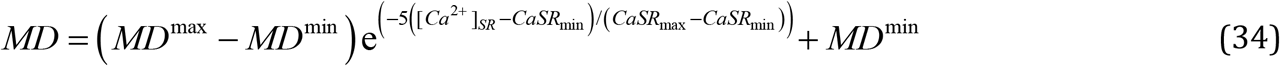

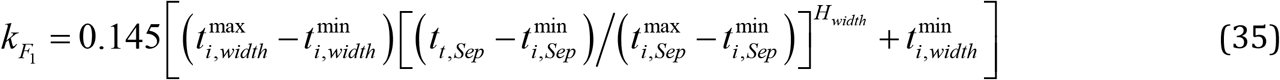

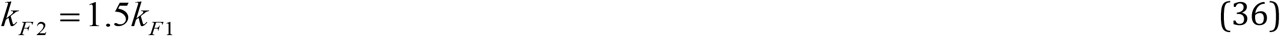

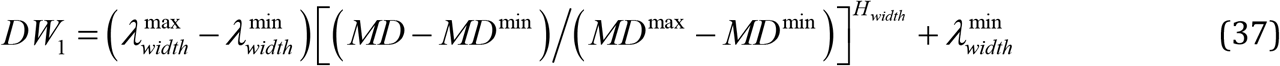

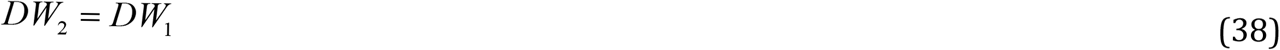

**Figure 8:**
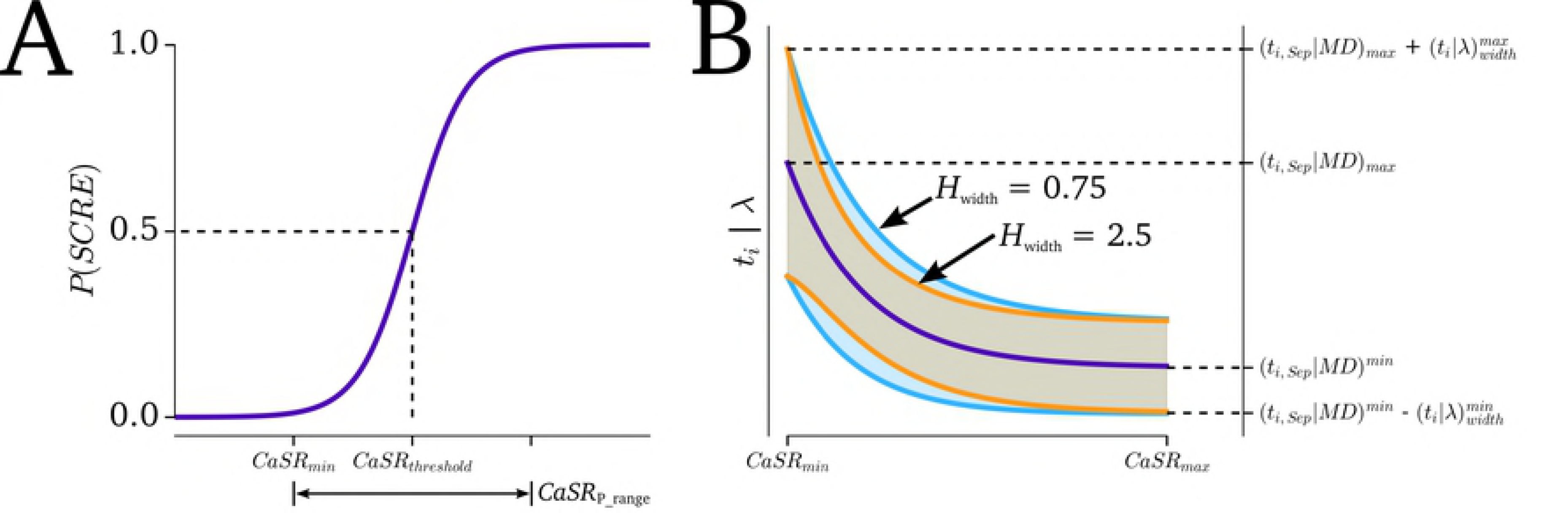
General Dynamic SRF model parameters. A - Illustration of the curve for probability of SCRE and its relation to the user defined parameters (*CaSR*_thresdhold_, *CaSR*_P_range_) and derived parameters (*CaSR*_min_). B - Illustration of the function form describing *t*_i,Sep_ and median duration (*MD*) (purple line) and its relation to *CaSR*_min_, *CaSR*_max_, *t*_i,sep_ ^min^, *t*_i,sep_ ^max^, *MD*^min^, *MD*^max^, and the how the range of the distributions varies with SR-Ca^2+^ (shaded regions) at two different non-linearity factors (*H*_width_ = 0.75, blue; = 2.5, orange).

## Results

In this section, the Dynamic Fit SRF approach for the three Ca^2+^ system conditions was validated by comparison of single cell SCRE in the 0D model with the 3D cell model. Generalisation of the approach is then demonstrated through: (i) parameterising the General Dynamic SRF model based on limited input data and (ii) integration of the SRF with an independent non-spatial cell model (and its native Ca^2+^ handling system). Finally, the potential applications of the models are illustrated through simulations of SCRE induced focal excitation in tissue models, demonstrating the emergence of different behaviour under different conditions.

### Validation of the Dynamic Fit Spontaneous Release Functions

The 0D model implementing the Dynamic Fit SRF model was first validated by comparison of whole-cell SCRE under Ca^2+^ clamp conditions with a second set of simulations (i.e., not those on which the model was derived) of the 3D cell model (Figure 9). These simulations highlight the strong agreement in both waveform variation (Figure 9A,B), the distributions (Figure 9C-D) and their summary properties (Figure 9E). Similar agreement was also observed for both of the remodelled conditions (Figure 10).

**Figure 9:**
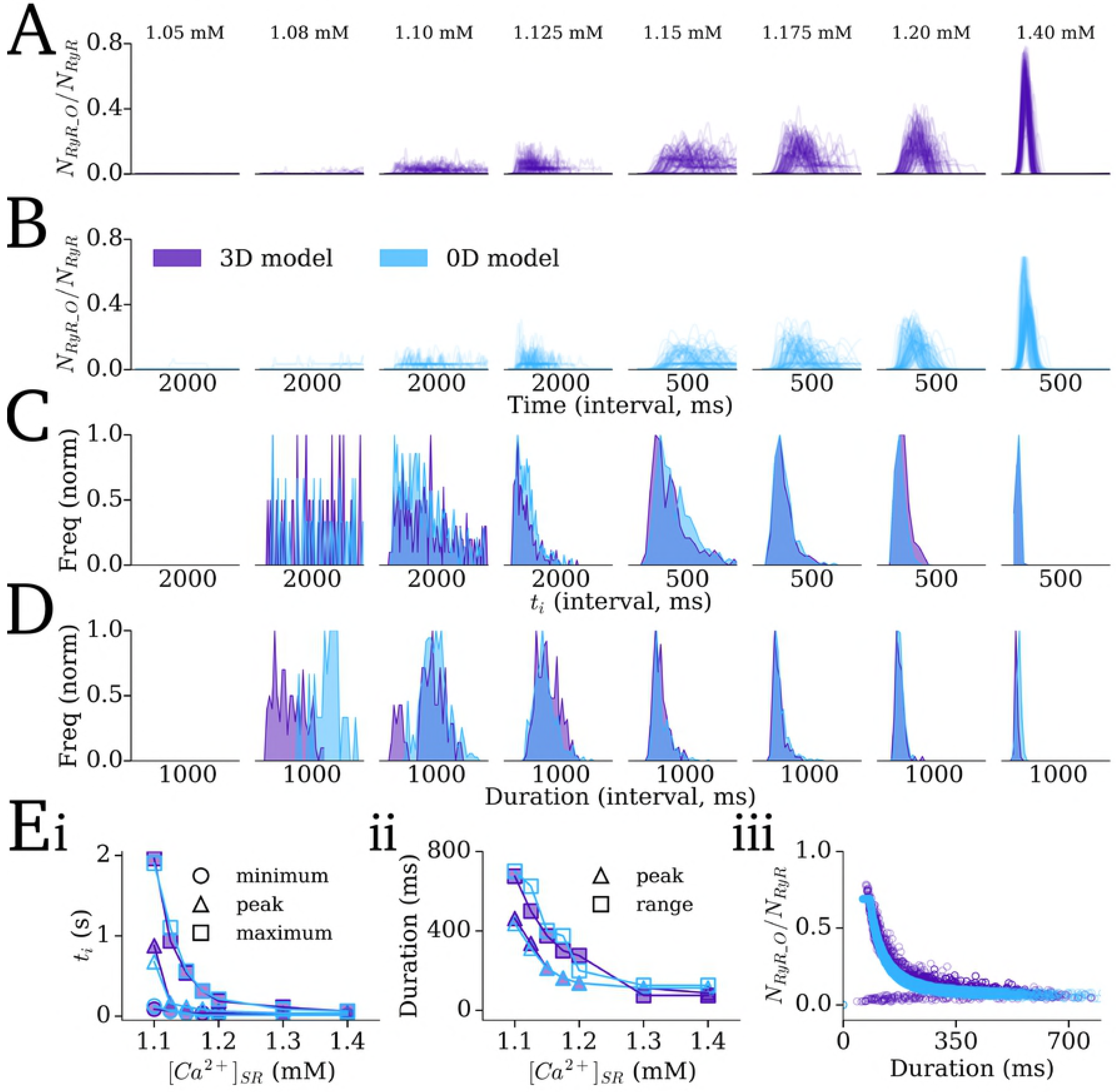
Validation of the control SRF through Ca^2+^ clamp. Results of 250 simulations at different SR-Ca^2+^ concentrations (labels on top of panel A) for the 3D model (purple) and 0D model with control Dynamic Fit SRF implementation (blue), showing: traces of the open RyR from 100 simulations at each SR-Ca^2+^ value (A,B); Histograms of the initiation time (C) and duration of the waveform (D); Summary of measured distribution parameters (Ei,ii), and scatter plot of peak open RyR against duration (Eiii). The values given on the *x*-axis of panels A-D refer to the total time interval of the plot, not absolute values.

**Figure 10:**
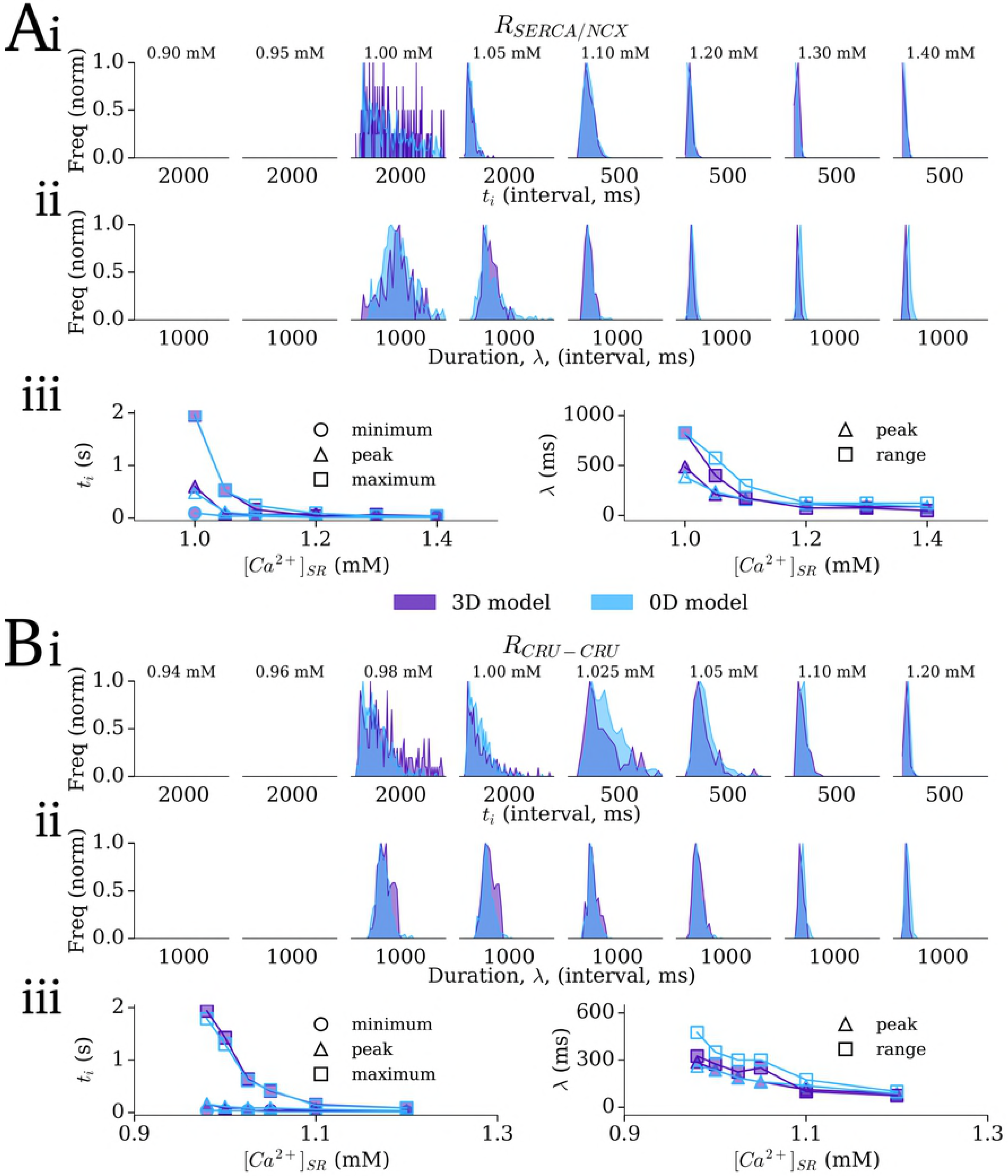
Validation of the remodelling SRF through Ca^2+^ clamp. Results of 250 simulations at different SR-Ca^2+^ concentrations (labels on top of panel A and B) for the 3D cell model (purple) and 0D model (blue) for the SERCA up-regulated/NCX down-regulated remodelling model (R_SERCA/NCX_, A) and increased CRU-CRU coupling remodelling model (R_CRU-CRU_, B). Histograms of the initiation time (i) and duration of the waveform (ii); Summary of measured distribution parameters (iii). The values given on the x-axis of panels (i-ii) refer to the total time interval of the plot, not absolute values.

Secondly, the 0D and 3D models were compared across the range of ionic models (see Methods: Action Potential and Tissue Models) and remodelling/ISO conditions (see Methods: Proarrhythmic conditions and analysis protocols). Examples of dynamics emerging in conditions close to the SR-Ca^2+^ threshold (Figure 11Aa) and high above it (Figure 11Ab) show good agreement between the 3D and 0D models (compare Figure 11Ai-ii with Aiv-v), importantly capturing the difference between the conditions. The distributions of initiation time and the probability of triggered APs across all conditions tested which resulted in notable SCRE also show good agreement (Figure 11B), confirming the ability of the 0D models to dynamically reproduce the SCRE of the 3D cell models and capture cellular and condition dependent differences.

**Figure 11:**
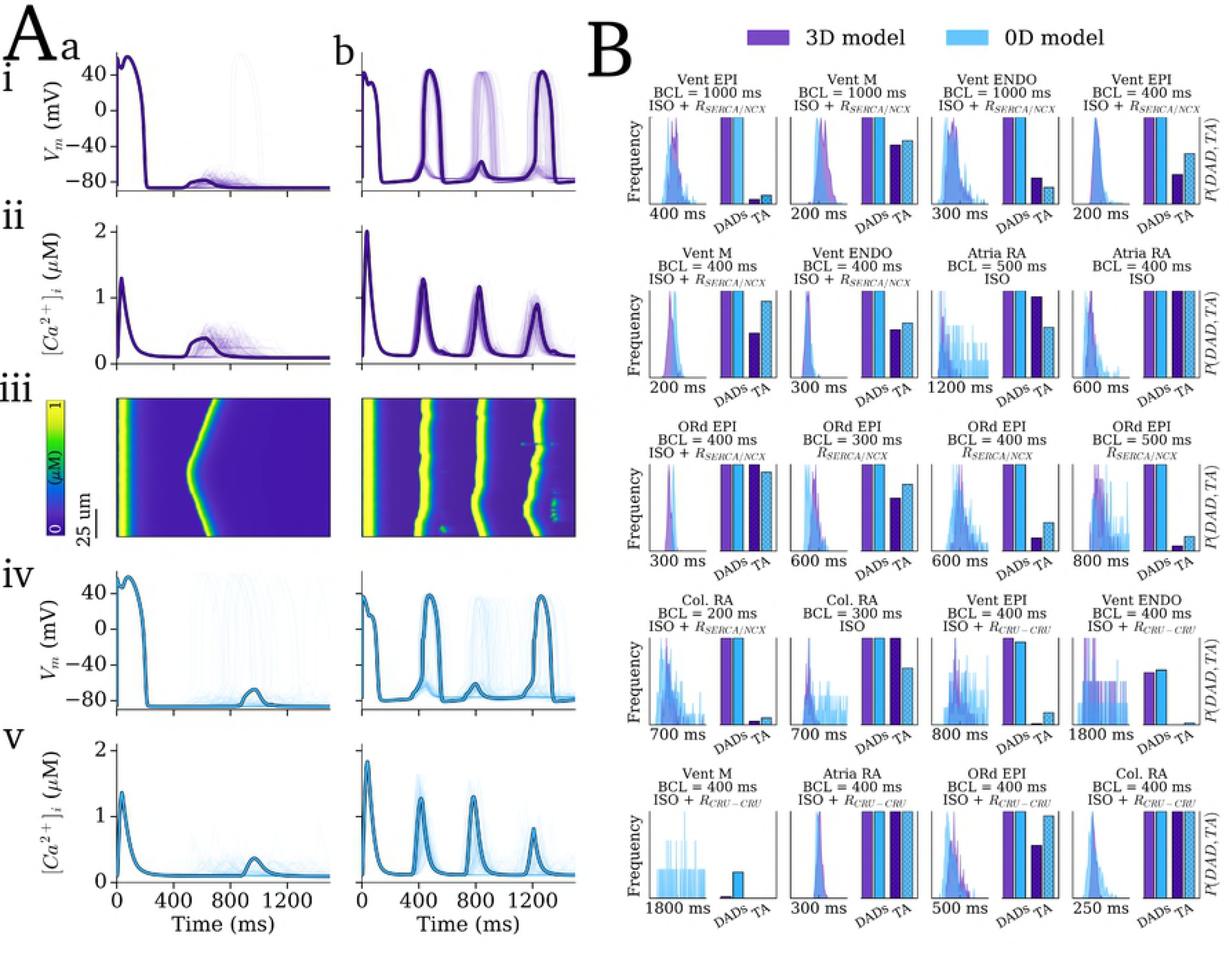
Validation of the SRF under dynamic pacing conditions. Results of 250 simulations for 20 different pacing conditions in which notable SCRE occurred. A - 100 examples (with one highlighted) of SCRE occurring in the 3D model (purple, i-iii) and 0D model (blue, iv-v) for two different conditions which resulted in SR-Ca^2+^ close to threshold (a, corresponding to Vent EPI, BCL = 400 ms, *R*_CRU-CRU_) and above it (b, corresponding to Vent ENDO, BCL = 400 ms, ISO + *R*_SERCA/NCX_). The linescans in (iii) correspond to the highlighted trace in (i-ii). Final paced beat and subsequent quiescent period is shown. B - Histograms of SCRE initiation time (left of each panel) and incidence of DADs and TA (bars, right of each panel) for the 20 different conditions (panel titles correspond to cell model, pre-pacing BCL and pro-SCRE conditions); the x-axis label for the histogram plots refers to the total range over which the plot is shown, rather than absolute values.

### Generalisation of the approaches

To demonstrate generalisation of the developed approaches, the SRF were first integrated with an independent cell model and its native Ca^2+^ handling system (Figure 12A). The human atrial AP model of Grandi et al., 2011 [51] was selected due to its detailed Ca^2+^ handling system containing a physiologically representative model of CICR and RyR model suitable for integration with SCRE waveforms. A General Dynamic implementation was parameterized to the SR-Ca^2+^ observed in that model in order to reproduce, for example, rate dependent susceptibility to SCRE (Figure 12A). This demonstrates the potential suitability for direct integration with available contemporary, non-spatial AP models, without the requirement to replace the native intracellular Ca^2+^ handling system.

Furthermore, approaches to directly incorporate experimental data were demonstrated. Starting with a suitable dataset, for example as found in the analysis of rate-dependence of SCRE in the rabbit atria presented in Workman et al., 2012 [52] (Figure 12Bi-ii), SCRE occurring at each pacing rate could be directly integrated (through the Direct Control implementation) to study the potential for observed SCRE to result in focal excitation. Moreover, by relating pacing rate with SR-Ca^2+^ in the model, the measured data could be plotted against model SR-Ca^2+^ and the General Dynamic implementation parameterized to fit these data (Figure 12Biii-iv). Parameterising the duration of SCRE from the data describing AP amplitude is non-trivial, partly due to the indirect experimental measure (i.e., not the peak of open RyRs), and partly due to the role of excitation determining this amplitude. Never-the-less, the duration distribution could be approximated, utilising the threshold for TA (dotted lines, Figure 12Bii, iv). Implementation of this General Dynamic model resulted in the expected rate dependence of the timing and amplitude of SCRE (Figure 12C).

**Figure 12:**
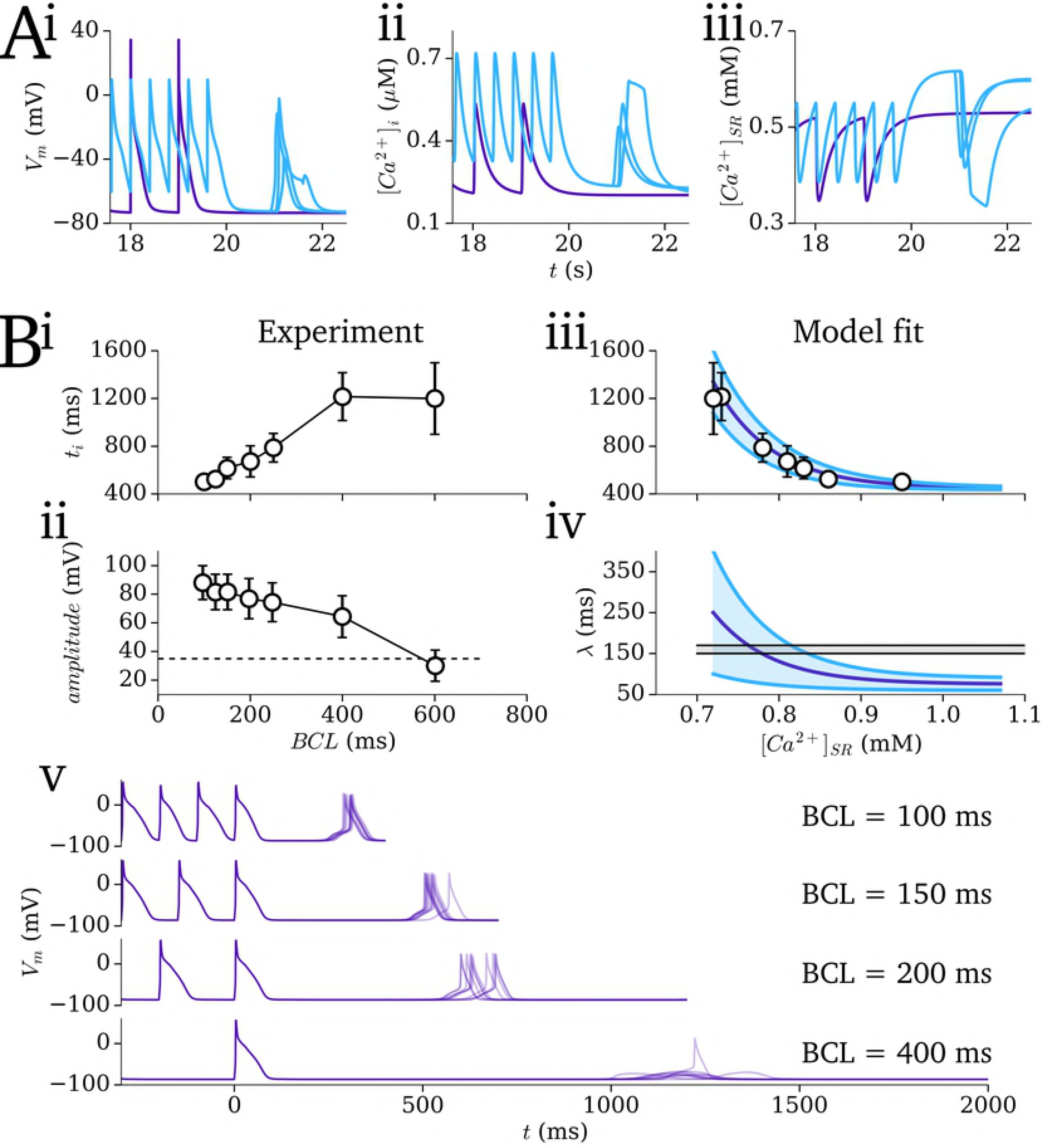
Generalisation of the approaches. A - Illustration of the General Dynamic SRF implementation integrated directly into the Grandi et al. 2011 human atrial model [51], showing membrane potential (i), intracellular Ca^2+^ (ii), and SR-Ca^2+^ associated with slow (BCL = 100 ms) and 3 simulations of rapid (BCL = 400ms) pacing. The threshold for SCRE was set to 0.6 mM. B - Parameterising the General Dynamic SRF to experimental data of Workman et al. 2012 [52]. (i) Initiation time and (ii) amplitude of SCRE at different cycle lengths in experiment; (iii) Fitting the initiation time and variability to the experimental data plotted against model SR-Ca2; (iv) approximating the duration of the waveform as a function of SR-Ca^2+^ utilising the durations (amplitudes) corresponding to the threshold for excitation (dotted lines). Parameter values for the General Dynamic SRF model for each case are provided in the Supplementary Material S1 Text (Model Description).

### Simulation at the tissue scale

The potential for the models to simulate SCRE at the tissue and organ scale was illustrated through pacing tissue models under an equivalent rapid pacing-quiescent protocol to the single cells (see Methods: Pro-arrhythmic conditions and analysis protocols). Under the right conditions (i.e., significant SR-Ca^2+^ loading and thus large-scale release events) a triggered action potential emerged from a single focus and propagated throughout the tissue, demonstrated in multiple tissue models (Figure 13). Multi-focal activations were also observed.

To compare the different cell model conditions, 2D sheets were pre-paced at cycle lengths of 200-1000 ms under all conditions (control, 2×remodelling, ±ISO). The different SCRE dynamics observed in single 3D cells under these conditions (Figure 11B) was accentuated in tissue, wherein focal excitations only emerged under conditions which resulted in significant TA (Figure 14). Note also the important impact of reduced *I*_K1_, present in the atrial cell models and *R*_SERCA_NCX_ remodelling conditions, on allowing the emergence of tissue focal excitation.

**Figure 13:**
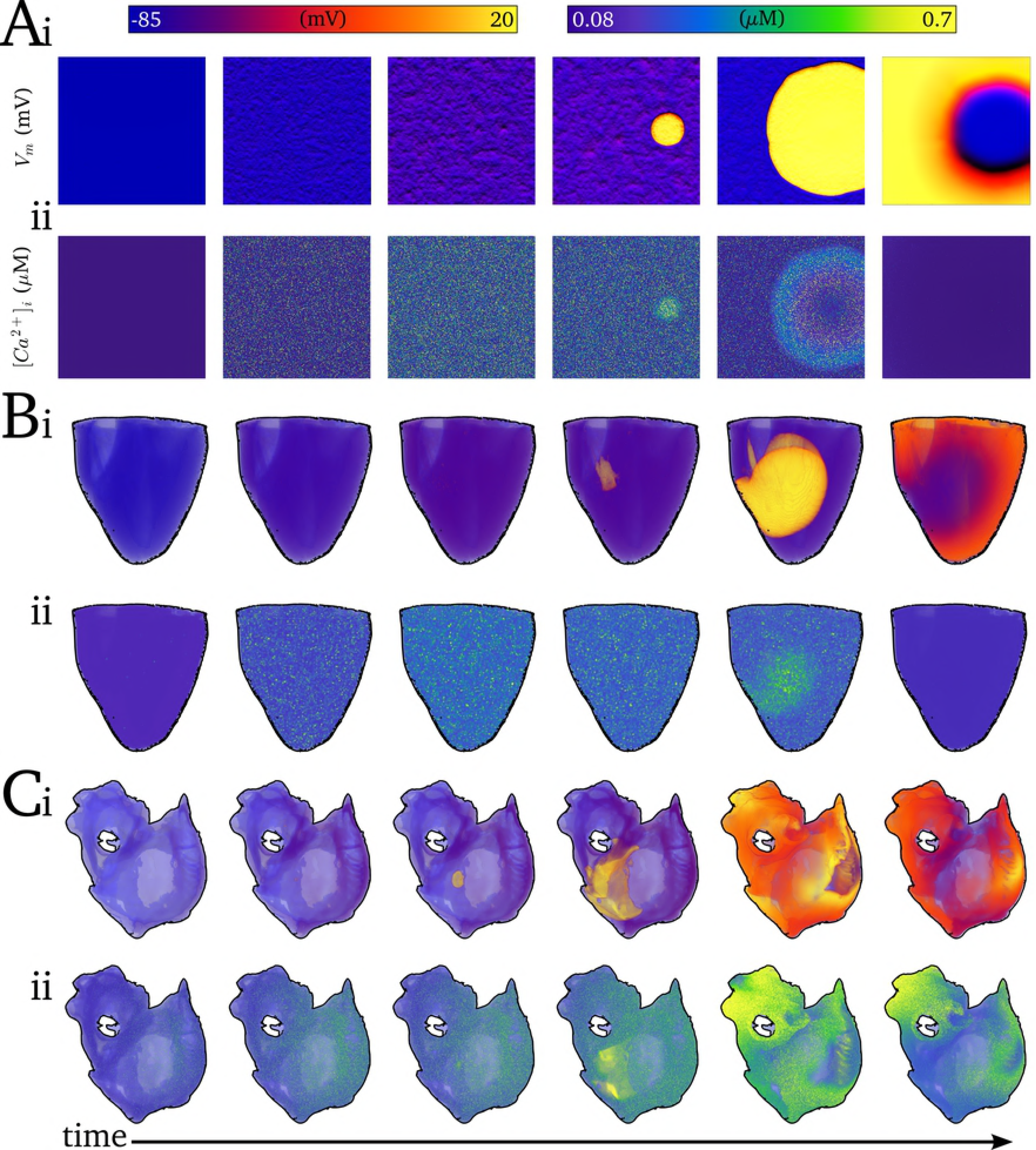
Development of SCRE mediated ectopic focal excitation. Snapshots of SCRE mediated focal excitation in idealised 2D sheet (A), whole-ventricle (B) and whole-atria (C) tissue models, showing the membrane potential (i, mV) and intracellular Ca^2+^ concentration (ii, μM).

**Figure 14:**
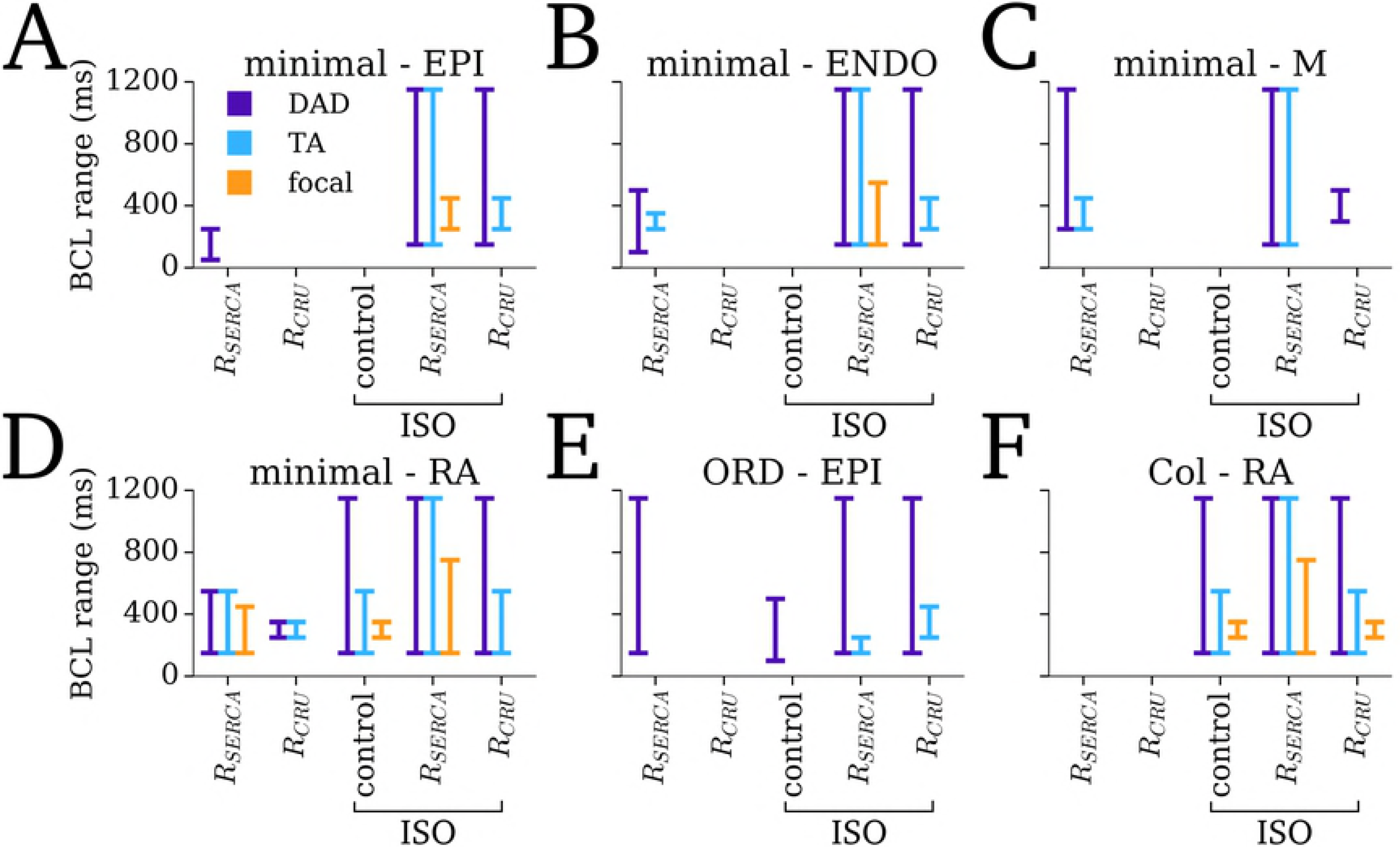
The emergence of SCRE at the tissue scale in different models. The BCL range over which activity corresponding to single cell DADs (purple) and TA (blue) and tissue focal excitation (orange) is shown for each of the different cell models/regions (A-F) under different conditions (*x-axis* labels remodelling, and control and remodelling + ISO). Note that no significant SCRE was observed for control conditions for any model, and so this condition has not been included in the figure.

## Discussion

### Summary

An increased incidence of spontaneous calcium release events (SCRE) is frequently observed in isolated cardiomyocytes from the diseased myocardium [6–10], and their pro-arrhythmic coupling to the membrane potential through activating inward NCX current has led to the hypothesis that these dysfunctional Ca^2+^ handling phenomena play a role in the initiation and dynamics of complex arrhythmia conduction patterns. However, investigating these multi-scale mechanisms presents a significant challenge, both for experimental and simulation approaches, and thus the precise mechanisms and potential importance of these events manifesting as tissue-scale arrhythmia have yet to be fully described.

In this study, a multi-scale computational approach was developed to simulate the dynamics of stochastic SCRE in organ-scale models of cardiac excitation. The computational framework comprises a hierarchy of models (Figure 2) encompassing the microscopic(3D cell model), mesoscopic(0D cell model) and macroscopic-scales (tissue models). Spontaneous Release Functions (SRF; Figure 4) were used to reproduce the morphology of SCRE in the 0D cell and tissue models (Figures 9-11, 13-14). Briefly, the waveform is fully described by its initiation time and duration, and randomly sampling these parameters from physiological distributions (Figure 7) ensured accurate and congruent stochastic variation in SCRE dynamics.

Multiple implementations were presented, providing the option to model SCRE either directly derived from the 3D cell model in specific, variable conditions, or determined by user input parameters. The potential for generalisation of the approaches developed was demonstrated through integration with an independent cell model and parameterisation based on experimental data (Figure 12). These models allow translation of single cell data (modelling or experimental) to the tissue scale and thus inference of arrhythmia mechanisms.

### Comparison to other approaches

Work from only a few research groups has attempted to simulate stochastic SCRE at the organ-scale [25,27,35,36,39]. These independent studies used broadly similar phenomenological approaches to overcome the inherent challenges of this multi-scale simulation as presented in this manuscript.

Further to providing an independent approach which forms a complementary tool to the previous models, which is of particular importance for theoretical investigation of systems with many unknowns and highly non-linear behaviour, the present study differs from those previously namely in: (i) the motivation to reproduce SCRE in tissue models in-line with that observed in specific (and variable) 3D cell models, for direct translation of single cell modelling studies to the tissue-scale; (ii) an approach which readily allows direct incorporation of experimental data; and (iii) the presentation of an open-source computational framework for congruent investigation of SCRE at single cell and tissue-scales with implementation designed for both direct translation and full controllability (Supplementary Material S1 Model Code).

## Limitations

The microscopic, 3D cell model, on which the computational framework is based, has inherent limitations: It has an idealised structure in which inter-dyad distances are constant throughout the cell volume and implements a compartmentalisation CRU approach, which ignores the complex underlying structures and their heterogeneity. A functional coupling sub-space was implemented to allow the propagation of Ca^2+^ waves in order to maintain this functionality with physiological Ca^2+^ transient magnitudes, as has been used in previous studies [9,53]. This was motivated by the desire to maintain SCRE associated *I*_NCX_ magnitude (dependent on Ca^2+^ transient amplitude) for accurate investigation of source-sink interactions. Incorporation of direct luminal regulation or sensitisation of the RyRs to drive propagation of Ca^2+^ waves, as has been used in other studies (e.g. [14]), may provide a more physiological solution. However, the primary purpose of the spatial cell model was to reproduce the dynamics, stochastic variability and SR-Ca^2+^ dependence of SCRE, and the model is therefore considered suitable for these ambitions.

The phenomenological approach presented provides an approximation to the 3D spatio-temporal dynamics and therefore does not capture the full complexity of the underlying behaviour. The method to derive the SRF in order to reproduce behaviour of the single cell model (i.e., not the general implementations) required a large volume of computationally intensive simulations to be performed (~ 5 000 hours of computation time per condition), although these do not need to be repeated once the parameters have been derived. Alternative approaches were also considered which have potential advantages, for example deriving a similar iterative-map approach to that presented in [24], parameterized to the spatial cell model dynamics, would require significantly less intensive simulations and perhaps provide a more robust underlying dynamic system. However, the analytical waveform approach presented also has advantages: based on whole-cell behaviour, parameter sets could be derived to describe underlying spatial models in limitless different conditions including sub-cellular heterogeneity and its variability, which may be significantly more challenging to reproduce with an underlying phenomenological dynamical system. The General Dynamic implementation also negates the requirement for simulations on which to derive the model, and permits direct parameterization to experimental data.

### Application of the framework

Due to the substantial detail required to sufficiently describe the approaches developed and their validation, only preliminary data were presented regarding modelling SCRE at the organ scale, illustrating a focal excitation and demonstrating the capability to reproduce and study cellular differences emerging at the tissue scale; More detailed application of the models to demonstrate the potential mechanisms of SCRE in arrhythmia is described in a related subsequent manuscript: Arrhythmia Mechanisms and Spontaneous Calcium Release: II From Calcium Spark to Re-entry and Back (Under Review). In this, the approaches are applied to: (i) describe the mechanism by which the independent cellular events manifest as a focal excitation; (ii) investigate the effect of cellular variability on the relationship between SR-Ca^2+^ and focal excitation; (iii) demonstrate two mechanisms by which SCRE can promote arrhythmic conduction patterns; and (iv) investigate the multi-scale interactions between SCRE and re-entrant excitation, revealing a functional mechanism for localisation of re-entrant scroll wave core and focal excitation.

It is intended that these approaches will be further developed and incorporated with sophisticated biophysically detailed cell models and experimentally validated simulation of cellular SCRE in multiple cardiac conditions, in order to suggest new experiments and contribute to detailed analysis of the role of SCRE in cardiac arrhythmia. Demonstration of the generalisation potential of the approaches is hoped to encourage those interested researchers in the community to integrate the presented framework with their cell models and simulation studies; open-source C++ code of the entire framework and detailed documentation is therefore provided in Supplementary Material S1 Model Code.

## Conclusions

The multi-scale cardiac modelling approaches described in this manuscript and accompanying model code present the possibility to model the impact of stochastic, sub-cellular calcium dynamics on organ scale arrhythmic excitation patterns with congruent detailed cellular and tissue simulations or parameterised to specific experimental datasets. The mechanistic insight gained from the application of these approaches may help to improve understanding and management of cardiac arrhythmia.

## Acknowledgements

I would like to thank Drs Al Benson and Izzy Jayasinghe, University of Leeds, for providing experimental data used in Figure 1, and also to Al for providing comments and feedback on the manuscript.

## Supplementary material

**S1 Text Model description:** Document containing full model equations and parameters for all of the cell models and SRF presented in this study.

**S1 Code:** C/C++ code containing all implementations presented in the study [Note: will be supplied at revision stage following any updates due to review].

**S1 Video Illustration of Ca^2+^ clamp and SCRE dynamics:** Video corresponds to snapshots shown in Figure 3. Left panels show proportion of open RyR (upper) and the intracellular (purple/left axis) and SR (blue/right axis) Ca^2+^ concentrations (lower). Right panel shows the intracellular Ca^2+^ concentration in the 3D volume of the idealised cell model. Video covers SR-Ca^2+^ below threshold, just above, and significantly above.

